# Extending Musculoskeletal Modelling Validation Beyond Gait: Knee Loading and Kinematics During Daily Activities

**DOI:** 10.64898/2026.10.24.761008

**Authors:** Enrico De Pieri, Sara C. Sarabadani Tafreshi, Sabrina Hörmann, Morten E. Lund, Philipp Damm, Pascal Schütz, William R. Taylor, Marc Bandi, Jörn Seebeck

## Abstract

Musculoskeletal (MSK) modelling enables non-invasive estimation of internal body loads, which could inform the design of novel orthopedic implants and guide personalized treatment strategies. Extensive validation is a pre-requisite for the translation of MSK-modelling frameworks into clinical applications. This study validates predicted knee loads and local kinematics across multiple activities of daily living (ADLs) using motion-capture data from 184 activity cycles collected in six subjects performing five ADLs from the CAMS-Knee datasets. Personalized models incorporated subject-specific anthropometric, radiographic, and implantation data, along with a 6-DOF elastic-foundation contact model between implant components and a surgically balanced ligament model derived from cadaveric testing. Across activities, predicted knee contact force magnitudes showed good agreement with instrumented implant measurements (RMSE 0.46 BW; R² = 0.81). The highest accuracy was found during level walking, whereas stair descent exhibited the largest discrepancies. Predicted implant kinematics showed qualitatively similar trends to fluoroscopic measurements across all ADLs, although with varying levels of accuracy across different degrees of freedom. EMG comparisons revealed broadly consistent activation timing between model predictions and experimental data, although gastrocnemius activation was poorly estimated during stair descent and downhill walking. By validating MSK model predictions across the complete range of activities available in the CAMS-Knee datasets, this study extends validation beyond previously analysed gait or bending tasks and reveals activity-specific discrepancies that reflect current limitations in modelling assumptions and would remain undetected in single-activity analyses. These findings provide a robust basis for future refinement of MSK models and support their development toward clinically meaningful applications.

## INTRODUCTION

Multibody musculoskeletal (MSK) modelling is a powerful tool for understanding movement biomechanics and estimating internal body loads in a non-invasive manner. Accurate knowledge of muscle and joint loading can support the clinical management of patients with motor impairments or musculoskeletal pathologies, guide the development of more effective treatment strategies, and inform the design and testing of novel orthopedic implants and surgical procedures [1–9].

Previous studies have measured *in vivo* joint loads using instrumented implants in patients undergoing total joint arthroplasty [4, 5]; however, these measurements remain available for only a limited number of patients and activities. While these measurements offer valuable insights, they are unlikely to fully represent the variability present in the broader population or across different loading conditions. MSK models can help overcome this limitation by estimating internal joint and muscle forces from externally measured kinematics and kinetics, which are readily obtained through modern motion capture systems. Through inverse dynamics, MSK models quantify the net joint moments required to perform a given movement. The redundancy of the musculoskeletal system is resolved by identifying an optimal combination of muscle forces that generates the required moments, enabling the computation of resultant joint contact forces [1, 10–12]. The in-silico nature of this approach further allows exploration of hypothetical scenarios to investigate how factors such as implant design, alignment, or surgical technique influence joint loading [13–16].

Despite their growing use in research [17, 18], MSK models have seen limited translation into clinical practice [19], product design, and surgical planning. This is largely due to a lack of extensive model validation [1, 2, 12, 19, 20]. Validation ensures that computational models appropriately represent reality [21] by assessing their ability to predict relevant variables with sufficient accuracy for the intended application [12, 22, 23]. In clinically relevant applications, establishing model credibility is paramount to patient safety [21]. Regulatory agencies are increasingly recognizing the potential of computational modelling in medical device development and are adapting testing and approval frameworks accordingly [2, 24–26]. Key components of any validation process include comparing model predictions against experimental data, defining the model’s range of validity, and determining the accuracy required for the intended (clinical) use [20, 22, 23, 27].

Direct and quantitative validation of MSK models is particularly valuable, as it builds confidence not only in the qualitative trends of the predictions but also in their quantitative accuracy [2, 22, 28]. Comparing predicted joint loads with those measured from instrumented implants offers a unique opportunity for such validation, especially when joint loads are the variables of interest. Because muscle forces are the primary contributors to internal joint loading [29], comparisons with instrumented implant data also provide an indirect validation of predicted muscle forces [12, 30].

Initiatives such as the “Grand Challenge Competition to Predict In Vivo Knee Loads” [12, 31] and the “Comprehensive Assessment of the Musculoskeletal System” (CAMS-Knee) [32] have provided benchmark datasets combining motion capture, surface electromyography (EMG), and in vivo joint contact forces, thereby enabling systematic testing of model performance. Previous direct validation studies have primarily focused on joint contact forces during level walking or sagittal-plane knee bending motions (e.g., squatting or sitting) [30, 31, 33–39]. However, these efforts offer only a partial view of model accuracy, as secondary joint kinematics are typically neglected [40]. In this regard, the CAMS-Knee datasets [32] currently represent the gold standard for MSK model validation, comprising data from six subjects fitted with instrumented knee implants, along with synchronized moving video-fluoroscopy recordings of local knee kinematics during multiple activities of daily living (ADLs), including level walking, downhill walking, stair descent, squatting, and chair rise.

This study therefore aims to develop an MSK modelling framework and assess the validity of its predictions against *in vivo* measurements from the CAMS-Knee datasets including knee contact forces and moments (KCF&Ms), EMG signals, and local knee kinematics during different ADLs.

## METHODS

### CAMS-Knee Datasets

The CAMS-Knee datasets [32] include experimental data from six patients (5 m, 1 f, age 68 ± 5 years, mass 88 ± 12 kg, height 173 ± 4 cm) who underwent total knee arthroplasty (4 left, 2 right), in which the tibial component of the cemented knee implant (INNEX FIXUC; Zimmer Biomet, Switzerland) was instrumented with strain gauges and a telemetry transmitter to measure 3D forces and moments of the tibiofemoral joint (90-100Hz, mean measurement error < 2% [41]). Whole-body kinematics and kinetics were collected using a 26-camera motion-capture system (Vicon Motion Systems Ltd., Oxford, United Kingdom) capturing at 100 Hz, synchronized with 6 force plates (Kistler Instrumentation, Winterthur, Switzerland) sampling at 2000 Hz. Seventy-five reflective skin markers were attached to predefined anatomical landmarks according to IfB [42], OSSCA [43], and Plug-in-Gait [44] marker sets. Furthermore, muscle activity for eight major lower limb muscles (medial and lateral vasti, rectus femoris, medial and lateral hamstrings, gastrocnemius medialis and lateralis, tibialis anterior) was measured bilaterally using a 16- channel wireless EMG system (Trigno, Delsys, USA) with signal delay of 48 ms. After the acquisition of a standing trial in an anatomic upright position, each subject completed a minimum of five valid trials for each of the investigated ADLs: level walking, downhill walking, stair descent, sitting and rising from a chair, and squat.

### Musculoskeletal Modelling

Motion capture data were used as input for a MSK modelling analysis in the AnyBody Modeling System (AMS; version 8.1.4; AnyBody Technology A/S, Aalborg, Denmark) [10] (Figure 1).

**Figure 1:**
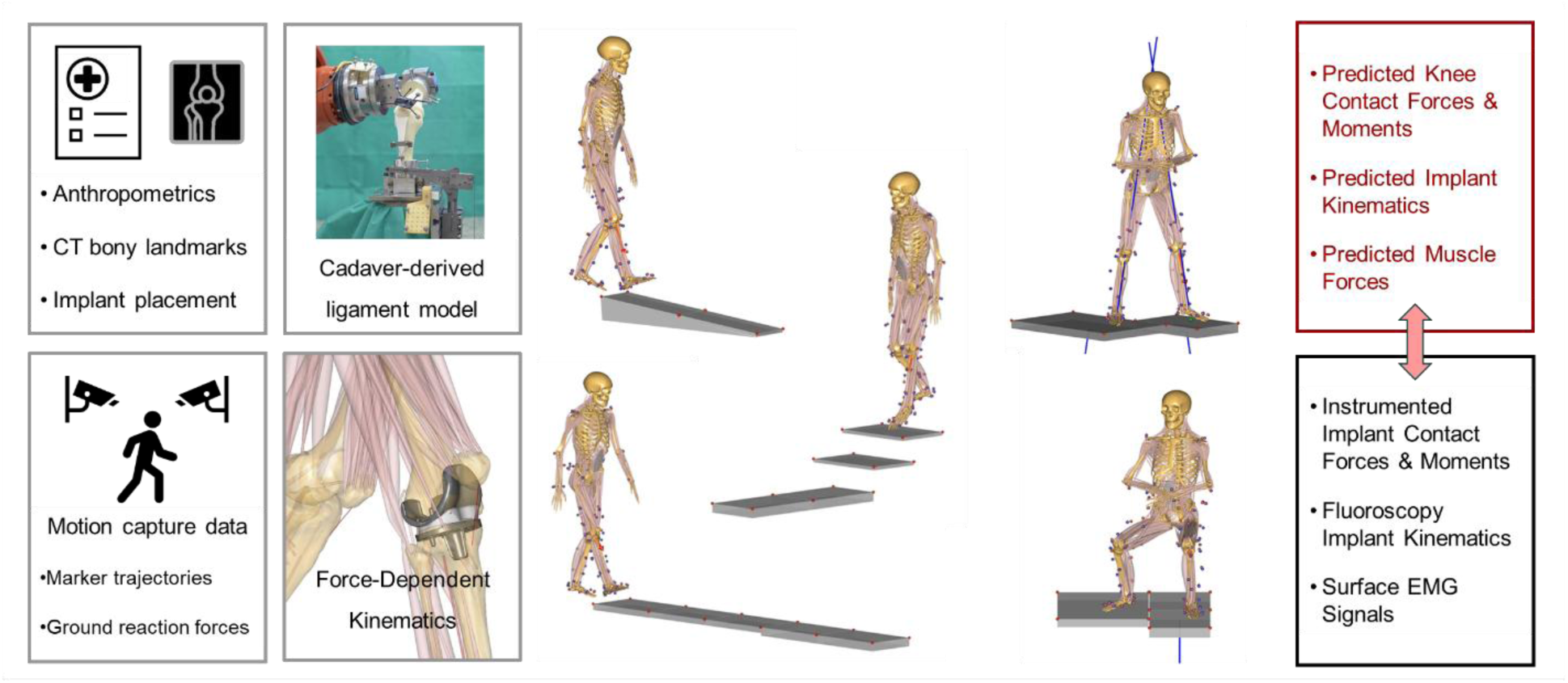
MSK modelling and validation workflow: personalized models incorporated available subject-specific anthropometric, radiographic, and implantation data, along with a 6-DOF FDK contact model between implant components and a surgically balanced ligament model derived from in-house cadaveric testing. Predicted knee contact forces and moments, local implant kinematics, and muscle forces during 5 ADLs (level walking, downhill walking, stair descent, sitting and rising from a chair, and squat) were compared against in vivo measurements from instrumented implants, fluoroscopy, and surface EMGs, respectively.

#### Model Scaling

A generic model, consisting of a simplified upper body (lumbar region, rigid trunk, neck, and head) and 11 segments representing the lower limbs (pelvis, right and left femurs, patellas, shanks, tali, and feet) was used. The musculoskeletal geometry of the lower limbs was based on the Twente Lower Extremity Model (TLEM) v2.2 (https://anyscript.org/ammr/body/leg_tlem2_model.html), derived from the cadaveric dataset TLEM 2.0 [45] and further refined for improved anatomical fidelity [28] (Appendix A available in the Supplemental Materials). The generic model was initially scaled to match each subject’s height and mass. Pelvic width, femoral length, and tibial length were then scaled to match the available CT-based bony landmarks and joint centers’ coordinates, while the remaining segmental dimensions were scaled from marker data collected during a static standing reference trial [46, 47]. Furthermore, tibial torsion was also computed from available CT-derived bony landmarks for the implanted leg as the rotation between the distal intermalleolar axis and the normal to the proximal AP axis of the tibial tray, which was assumed to be aligned with the axis connecting the posterior cruciate ligament (PCL) attachment point to the tibial tuberosity (TT). The modelled tibial geometry was morphed with a 3D radial-basis-function transformation to match the subject-specific torsional value computed from the CT data by including a transversal rotation between the tibial proximal and distal sections, which was distributed along the tibial shaft [48].

The model contains 55 muscle actuators for each leg, discretized into 169 fascicles in accordance with the original TLEM dataset [45, 49]. The maximal strength of each muscle was scaled according to subject-specific mass and estimated fat composition [50, 51]. The muscle fascicles were modelled as constant strength actuators, therefore neglecting force-length and force-velocity relationships. Different gastrocnemius and soleus muscle models were evaluated and discussed in Appendix B available in the Supplemental Materials.

#### Virtual Implantation

Subject-specific positioning of both femoral and tibial implant components in the MSK models was achieved through the in-house software ZiBRA™ Anatomical Modeling System (Zimmer Biomet, Switzerland) [52]. Starting from the scaled and morphed TLEM geometries of femur and tibia, the virtual implantation followed the surgical guidelines for the INNEX FIXUC system. Subject-specific implant sizing and orientation relative to known bony landmarks from the CAMS-knee dataset were used to inform the virtual implantation. Modelled flexion, coronal, and external rotation alignment of the femoral component, as well as the coronal alignment and posterior slope of the tibial tray, matched the orientation data from the CT scans. As no proximal tibial bony landmarks’ coordinates were available in the dataset, the AP axis of the tibial tray was aligned with the PCL-TT axis, following standard surgical guidelines for this specific implant. Resection depth, anterior-posterior and medial-lateral referencing for both components were adjusted to match the articular surfaces of the scaled bone geometries, according to implant-specific surgical guidelines.

The instrumented-tibial-tray-based coordinate system (CS_JWI_TIB_, [53]), in which KCF&Ms were measured, was reproduced in the MSK models. Furthermore, specific reference frames for the analysis of secondary joint kinematics were defined in the femoral and tibial components. The femoral component kinematic (FCkin) reference frame was centered in the midpoint between medial and lateral flexion facet centers, with the X axis pointing anteriorly, the Y axis proximally, and the Z axis to right. The medial and lateral flexion facet centers were derived from the axis of a cone fitted to the posterior implant condyles. The tibial baseplate kinematic (TBkin) reference frame was centered in the midpoint between the medial and lateral dwell points of the articular surface, with the X axis pointing anteriorly, the Y axis proximally, and the Z axis to right.

#### Joint Modelling

The hip was modeled as a 3-degrees of freedom (DOF) ball-and-socket joint, while talocrural and subtalar joints were modeled as 1-DOF hinges. Additionally, the position of the patella was defined as a function of the knee flexion angle [45].

The tibiofemoral joint was defined as a 6-DOF joint using the force-dependent kinematics (FDK) approach [30, 54], in which knee flexion was driven by marker tracking while secondary tibiofemoral kinematics were computed by solving the instantaneous quasi-static equilibrium between contact forces, external loads, muscle forces, and ligament constraints. Elastic foundation contact models were used to describe the contact forces between femoral component and tibial inlay, separately for the medial and lateral compartments, with a pressure module of 10 GN/m^3^ [30, 54, 55].

#### Ligament Modelling

Due to the lack of intra-operative ligament assessments for the CAMS-Knee subjects, a numerical model for the behaviour of the knee ligaments was instead derived from in-house robotic cadaveric experiments. The laxity profiles of a series of cadaveric specimens were evaluated under specific loading conditions on a 6-DOF KUKA robotic simulator (KR140 comp; KUKA, Augsburg, Germany) equipped with a 6 DOF force-torque sensor (ATI, Apex, USA) [56]. Through subsequent resection of individual ligaments and soft tissue structures, and the replay of previously measured knee kinematics with the robotic arm, the kinetic contribution of each ligamentous structure was differentially computed, allowing the elastic response to be modelled at any given tibiofemoral position. All the individual ligament models were then used to create a synthetic 6-DOF numerical model of the overall joint elasticity for each cadaveric specimen [57]. The joint elasticity function (JEF) of a single cadaveric specimen was then adapted to the individual CAMS-Knee subjects, so that it would present the same laxity behaviour of the INNEX- implanted cadavers while respecting the additional constraints of CAMS-subject-specific bone dimensions and CT-derived implant positioning. This approach preserved the internal consistency of the experimentally derived ligament model without requiring assumptions regarding individual ligament slack lengths or operating ranges. Specifically, all the JEFs adapted to the CAMS-subjects presented varus-valgus laxity profiles qualitatively similar to the series of INNEX- implanted cadaver specimens throughout the range of knee flexion. In particular, the laxity profiles in full extension and at 90° of knee flexion are clinically relevant, as they correspond to the gap-balancing assessment typically performed intra-operatively by surgeons, indicating that the JEFs incorporated into the MSK models reproduced surgically balanced knees.

#### Marker Tracking and Inverse Dynamics

The laboratory force plate set-up was recreated in the simulations following the description included in the CAMS-Knee datasets. The marker trajectories and GRF data were filtered through a 2^nd^ order low-pass Butterworth filter with cut-off frequencies of 6 and 12 Hz, respectively. A kinematic analysis based on marker trajectories was conducted to compute primary joint kinematics [46, 47]. Secondly, an inverse dynamics analysis, based on a 3^rd^ order polynomial muscle recruitment criterion, was performed to calculate required muscle activations and forces, resulting joint moments and contact forces, as well as secondary tibiofemoral kinematics through FDK [11]. As polynomial muscle recruitment criteria are sensitive to the uneven discretization of anatomical muscles into smaller fascicles, a muscle-volume-based normalization factor was included in the muscle recruitment cost function [30, 58–60].

### Data Analysis

The modelling and validation workflow was automated through the workflow management system Snakemake [61] and the toolkit AnyPyTools [62] in the Python programming language (version 3.11, Python Software Foundation, https://www.python.org/).

Both measured and predicted data were cropped into valid activity-specific cycles. For level walking, downhill walking, and stair descent, cycles were time-normalized from heel strike (0%) to heel strike (100%) and interpolated to 1% increments (101 points). For squatting, a complete cycle was defined from upright standing (0%) to deep flexion (50%) back to upright (100%). Initiation and completion of the activity were set as the knee flexion angle velocity crossed a 10°/s threshold, while the cycle mid-point was defined when knee flexion angle reached its maximum value. For sit-to-stand-to-sit, initiation (0%) and completion (100%) of the cycle were identified when the velocity of the body center of mass crossed a threshold of 0.03 m/s, while the standing phase (45-55%, in between standing up from the chair and sitting back down on it) was approximately centered around the minimum knee flexion angle and delimited by the knee flexion angle velocity crossing a threshold of 20°/s.

The analysis was limited to the limb with the instrumented TKA, for a total of 184 activity cycles with valid kinetic data across the five ADLs and 6 subjects. Measured and predicted KCF&Ms were reported in CS_JWI_TIB_. KCFs were normalized by body weight (BW), while KCMs were normalized by body mass (BM). Relative translations and rotations of the femoral component (FCkin) with respect to the tibial baseplate (TBkin) (Cardan rotational sequence: Y-X-Z) were computed for both MSK models and video-fluoroscopy data in 156 activity cycles with both valid kinetic and fluoroscopy data.

Root mean square error (RMSE) and coefficient of determination (R^2^) were computed between measured and predicted KCF&Ms and secondary kinematics across patients and activities. The maximum errors between predictions and measurements per trial are also reported.

Furthermore, the predicted muscle forces were qualitatively compared against experimentally measured EMG signals for each trial. The raw EMG signals were bandpass-filtered (2^nd^ order Butterworth filter >20 Hz, < 500 Hz), rectified, and low-pass filtered (2^nd^ order Butterworth <6 Hz). Trials in which EMG signals visually presented motion artefacts were manually excluded from this analysis. Valid EMG signals were normalized to the maximum value recorded across all trials and activities for each subject [35], with left and right limbs normalized independently. Predicted muscle forces were defined as the sum of the forces generated by all fascicles constituting each muscle in the MSK models. Each muscle force was also normalized to its maximum value across all trials and activities for each subject.

## RESULTS

### KCF&Ms

Across all activities, the predicted KCF magnitude presented good agreement with the data from the instrumented implants (Figure 2), with a RMSE of 0.46 BW [inter-subject range: 0.35 – 0.61 BW] and a R^2^ of 0.81 [0.57 – 0.91] (Table 1). The proximal KCF component presented a similar level of agreement (RMSE = 0.45 BW, R^2^ = 0.82), while the anterior and the lateral component showed poor agreement (anterior: RMSE = 0.22 BW, R^2^ = -2.02; lateral: RMSE = 0.08 BW, R^2^ = - 0.42). The KCM showed moderate to good agreement in terms of magnitude (RMSE = 0.08 Nm/kg and R^2^ = 0.7) and adduction component (RMSE = 0.07 Nm/kg and R^2^ = 0.8) and weak to poor agreement for the internal rotation (RMSE = 0.03 Nm/kg and R^2^ = 0.23) and extension (RMSE = 0.12 Nm/kg and R^2^ = -0.83) components. When comparing the individual activities, the models performed the best during level walking (KCF magnitude: RMSE = 0.29 BW, R^2^ = 0.92) and the worst during stair descent (KCF magnitude: RMSE = 0.70 BW, R^2^ = 0.69). Stair descent was also characterized by the largest maximum error (KCF magnitude: 1.76 BW, mean value computed across the maximum errors of all trials of all subjects).

**Figure 2:**
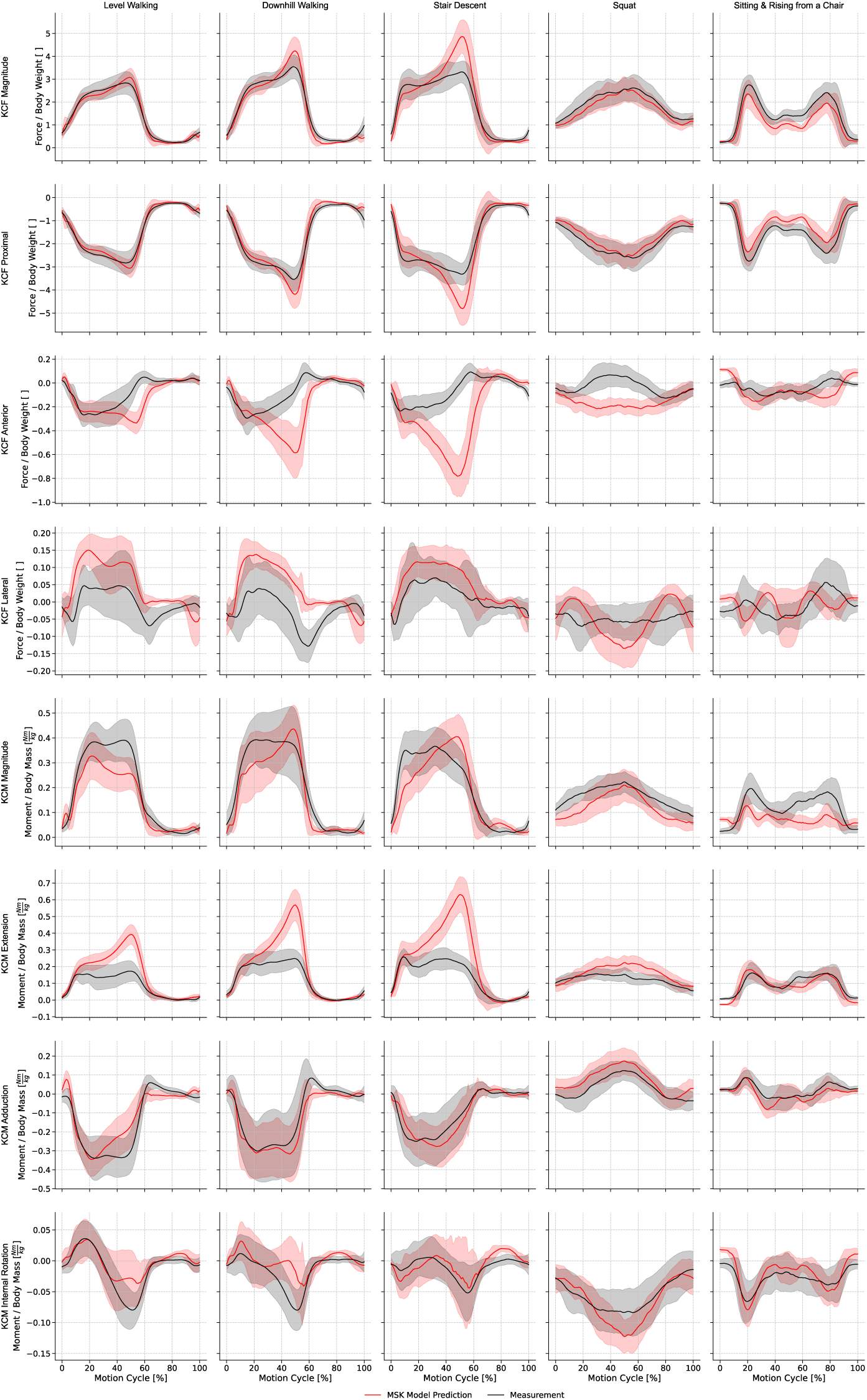
Mean (± SD) predicted knee contact forces and moments (KCF&Ms; red) compared with experimental measurements (black) from the CAMS-Knee datasets [32] during different ADLs. Force components include KCF magnitude, proximal (+)–distal (−), anterior (+)–posterior (−), and medial (+)–lateral (−) directions. Moment components include KCM magnitude, extension (+)– flexion (−), adduction (+)–abduction (−), and internal (+)–external (−) rotation.

**Table 1:** Root mean square error (RMSE), coefficient of determination (R²), and maximum absolute error computed between measured and predicted KCF&Ms. RMSE and R² were computed across all available activity cycles of all subjects for the indicated activities. The maximum absolute error was computed for each activity cycle and then averaged across all relevant cycles for the indicated activities. Subject-specific RMSE, R², and maximum absolute error were also computed, with the minimum and maximum subject-specific values reported in brackets.

|  |  | All Activities | Level Walking | Downhill Walking | Stair Descent | Squat | Sitting and Rising<br>from a Chair |
| --- | --- | --- | --- | --- | --- | --- | --- |
| KCF Magnitude<br>[BW] | RMSE | 0.46<br>[0.35 - 0.61] | 0.29<br>[0.21 - 0.37] | 0.41<br>[0.25 - 0.55] | 0.7<br>[0.49 - 0.98] | 0.44<br>[0.2 - 0.88] | 0.46<br>[0.21 - 0.7] |
| | $R^2$ | 0.81<br>[0.57 - 0.91] | 0.92<br>[0.88 - 0.97] | 0.89<br>[0.8 - 0.97] | 0.69<br>[-0.37 - 0.86] | 0.58<br>[-0.62 - 0.94] | 0.7<br>[0.3 - 0.93] |
|  | Max error | 0.95<br>[0.76 - 1.18] | 0.67<br>[0.53 - 0.86] | 1.04<br>[0.52 - 1.35] | 1.76<br>[1.23 - 2.24] | 0.68<br>[0.4 - 1.34] | 0.86<br>[0.6 - 1.22] |
| KCF Proximal<br>[BW] | RMSE | 0.45<br>[0.35 - 0.6] | 0.29<br>[0.2 - 0.36] | 0.4<br>[0.25 - 0.53] | 0.68<br>[0.48 - 0.93] | 0.44<br>[0.2 - 0.88] | 0.46<br>[0.21 - 0.7] |
| | $R^2$ | 0.82<br>[0.61 - 0.91] | 0.93<br>[0.88 - 0.97] | 0.9<br>[0.81 - 0.97] | 0.71<br>[-0.22 - 0.87] | 0.58<br>[-0.62 - 0.95] | 0.7<br>[0.29 - 0.92] |
|  | Max error | 0.93<br>[0.76 - 1.16] | 0.66<br>[0.51 - 0.85] | 1.01<br>[0.53 - 1.34] | 1.7<br>[1.19 - 2.12] | 0.68<br>[0.4 - 1.36] | 0.86<br>[0.6 - 1.23] |
| KCF Anterior<br>[BW] | RMSE | 0.22<br>[0.16 - 0.28] | 0.16<br>[0.1 - 0.19] | 0.24<br>[0.16 - 0.35] | 0.37<br>[0.25 - 0.53] | 0.19<br>[0.11 - 0.3] | 0.11<br>[0.07 - 0.14] |
| | $R^2$ | -2.02<br>[-14.32 - -0.32] | -0.18 [-3.4 - 0.34] | -1.9 [-13.45 - -0.3] | -6.56 [-55.39 - -<br>1.91] | -2.82 [-17.94 - -<br>2.41] | -1.43 [-4.58 -<br>0.26] |
|  | Max error | 0.47<br>[0.4 - 0.52] | 0.4<br>[0.28 - 0.47] | 0.63<br>[0.48 - 0.76] | 0.87<br>[0.7 - 1.12] | 0.32<br>[0.18 - 0.56] | 0.23<br>[0.15 - 0.32] |
| KCF Lateral<br>[BW] | RMSE | 0.08<br>[0.06 - 0.11] | 0.09<br>[0.07 - 0.13] | 0.1<br>[0.07 - 0.11] | 0.07<br>[0.05 - 0.13] | 0.09<br>[0.04 - 0.13] | 0.07<br>[0.04 - 0.1] |
| | $R^2$ | -0.42<br>[-3.48 - 0.55] | -0.54<br>[-8.09 - 0.57] | -0.72<br>[-9.76 - 0.3] | 0.02<br>[-4.22 - 0.75] | -1.69<br>[-14.79 - -0.11] | -0.37<br>[-1.63 - -0.34] |
|  | Max error | 0.17<br>[0.13 - 0.21] | 0.2<br>[0.13 - 0.28] | 0.2<br>[0.13 - 0.29] | 0.15<br>[0.11 - 0.24] | 0.16<br>[0.08 - 0.25] | 0.15<br>[0.09 - 0.23] |
| KCM<br>Magnitude<br>[Nm/kg] | RMSE | 0.08<br>[0.06 - 0.09] | 0.08<br>[0.04 - 0.1] | 0.07<br>[0.06 - 0.1] | 0.09<br>[0.06 - 0.12] | 0.06<br>[0.03 - 0.08] | 0.08<br>[0.05 - 0.1] |
| | $R^2$ | 0.7<br>[0.48 - 0.85] | 0.75<br>[0.42 - 0.9] | 0.83<br>[0.35 - 0.94] | 0.67<br>[0.43 - 0.85] | -0.24<br>[-6.74 - 0.77] | -0.21<br>[-1.49 - 0.31] |
|  | Max error | 0.17<br>[0.14 - 0.2] | 0.17<br>[0.09 - 0.22] | 0.17<br>[0.13 - 0.24] | 0.23<br>[0.19 - 0.3] | 0.11<br>[0.07 - 0.15] | 0.15<br>[0.11 - 0.19] |
| KCM Extension<br>[Nm/kg] | RMSE | 0.12<br>[0.08 - 0.14] | 0.11<br>[0.07 - 0.16] | 0.13<br>[0.07 - 0.17] | 0.19<br>[0.14 - 0.23] | 0.07<br>[0.03 - 0.11] | 0.05<br>[0.04 - 0.06] |
|  | R<sup>2</sup> | -0.83<br>[-2.78 - 0.26] | -0.85<br>[-12.47 - 0.4] | -0.54<br>[-1.67 - 0.68] | -1.73<br>[-4.08 - -0.4] | -1.5<br>[-7.62 - 0.69] | 0.5<br>[-0.29 - 0.76] |
|  | Max error | 0.25<br>[0.18 - 0.3] | 0.24<br>[0.14 - 0.33] | 0.35<br>[0.22 - 0.44] | 0.45<br>[0.39 - 0.56] | 0.11<br>[0.07 - 0.17] | 0.11<br>[0.09 - 0.14] |
| KCM Adduction<br>[Nm/kg] | RMSE | 0.07<br>[0.04 - 0.1] | 0.09<br>[0.04 - 0.13] | 0.07<br>[0.05 - 0.1] | 0.07<br>[0.05 - 0.1] | 0.06<br>[0.03 - 0.1] | 0.04<br>[0.04 - 0.05] |
|  | R<sup>2</sup> | 0.8<br>[0.32 - 0.92] | 0.75<br>[0.17 - 0.91] | 0.88<br>[0.14 - 0.96] | 0.75<br>[-1.04 - 0.92] | 0.42<br>[-2.09 - 0.71] | 0.39<br>[-0.01 - 0.51] |
|  | Max error | 0.15<br>[0.11 - 0.19] | 0.18<br>[0.1 - 0.26] | 0.15<br>[0.1 - 0.21] | 0.19<br>[0.11 - 0.25] | 0.12<br>[0.06 - 0.16] | 0.09<br>[0.08 - 0.11] |
| KCM Internal Rotation<br>[Nm/kg] | RMSE | 0.03<br>[0.02 - 0.04] | 0.04<br>[0.02 - 0.04] | 0.04<br>[0.04 - 0.05] | 0.04<br>[0.02 - 0.04] | 0.04<br>[0.02 - 0.05] | 0.02<br>[0.02 - 0.03] |
|  | R<sup>2</sup> | 0.23<br>[-0.23 - 0.49] | 0.22<br>[-0.35 - 0.71] | -0.43<br>[-1.15 - 0.18] | -0.39<br>[-11.76 - 0.22] | 0.21<br>[-3.1 - 0.3] | 0.38<br>[-1.18 - 0.62] |
|  | Max error | 0.08<br>[0.05 - 0.09] | 0.08<br>[0.05 - 0.11] | 0.11<br>[0.1 - 0.13] | 0.09<br>[0.06 - 0.13] | 0.06<br>[0.04 - 0.08] | 0.05<br>[0.03 - 0.06] |

Results for the individual patients, including the contribution of the JEF to the KCF&Ms, are reported in Appendix C available in the Supplemental Materials.

### Implant Kinematics

The predicted local implant kinematics and fluoroscopic measurements also presented qualitatively similar trends during all ADLs (Figure 3). With regards to the individual degrees of freedom, the knee flexion angle showed good agreement with the experimental data (RMSE=5.01°, R^2^ = 0.97), the anterior and the proximal translations showed moderate agreement (RMSE=2.38mm, R^2^ = 0.75 and RMSE=0.72mm, R^2^ = 0.72, respectively), while the lateral translation, internal rotation and valgus angle presented poor agreement (RMSE=1.05mm, R^2^ = - 0.65; RMSE=5.09°, R^2^ = -1.35; and RMSE=1.22°, R^2^ = -1.71, respectively) (Table 2). During sitting and rising from a chair, the local knee kinematics differed from the experimental data when the subjects were still sitting or partially sitting (approximately 0-15% and 85-100% of the activity cycle), indicating that the FDK approach might struggle to find the correct local equilibrium point in a low-loading situation, but presented good agreement during the remainder of the motion task.

**Figure 3:**
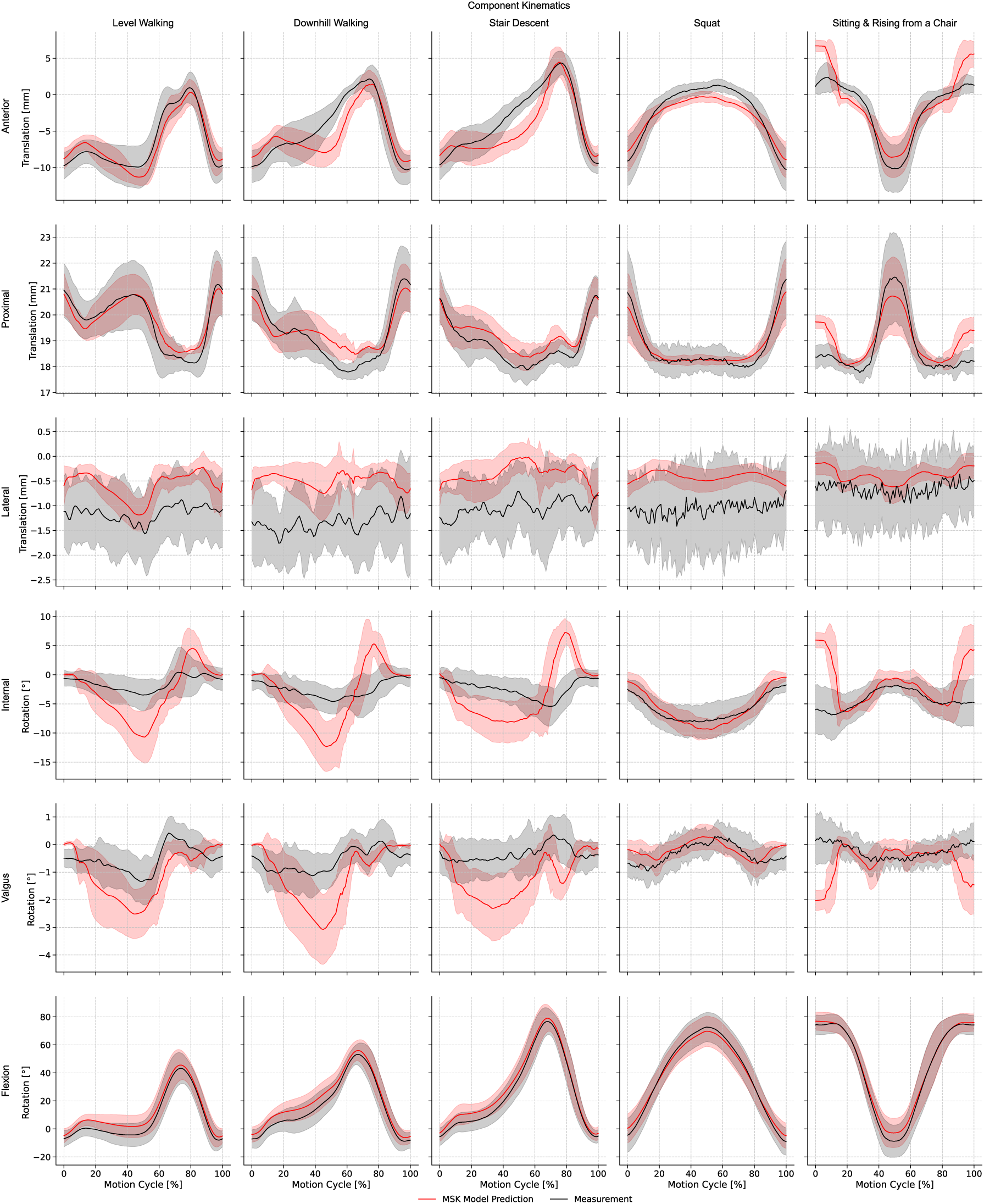
Mean (± SD) predicted implant kinematics (red) compared with experimental measurements (black) from the CAMS-Knee datasets [32] during different ADLs. Relative translations and rotations of the femoral component with respect to the tibial baseplate are shown. Translational components include anterior (+)–posterior (−), proximal (+)–distal (−), and lateral (+)–medial (−) directions. Rotational components include internal (+)–external (−), valgus (+)–varus (−), and flexion (+)–extension (−) rotations.

**Table 2:** Root mean square error (RMSE), coefficient of determination (R²), and maximum absolute error computed between measured and predicted implant kinematics. RMSE and R² were computed across all available activity cycles of all subjects for the indicated activities. The maximum absolute error was computed for each activity cycle and then averaged across all relevant cycles for the indicated activities. Subject-specific RMSE, R², and maximum absolute error were also computed, with the minimum and maximum subject-specific values reported in brackets.

|  |  | All Activities | Level Walking | Downhill Walking | Stair Descent | Squat | Sitting and Rising<br>from a Chair |
| --- | --- | --- | --- | --- | --- | --- | --- |
| Anterior<br>Translation<br>[mm] | RMSE | 2.38<br>[1.71 - 2.78] | 2.06<br>[1.13 - 2.59] | 2.8<br>[2.3 - 3.27] | 2.7<br>[1.95 - 3.5] | 1.66<br>[0.87 - 2.11] | 2.72<br>[2.23 - 3.45] |
| | $R^2$ | 0.75<br>[0.32 - 0.85] | 0.78<br>[0.41 - 0.91] | 0.62<br>[0.06 - 0.78] | 0.67<br>[0.11 - 0.84] | 0.82<br>[0.23 - 0.95] | 0.65<br>[-1.08 - 0.85] |
|  | Max error | 5.01<br>[4.08 - 5.49] | 4.75<br>[3.72 - 6.2] | 6.12<br>[4.71 - 6.99] | 5.48<br>[3.78 - 7.36] | 2.98<br>[1.72 - 3.68] | 6.2<br>[5.38 - 7.22] |
| Proximal<br>Translation<br>[mm] | RMSE | 0.72<br>[0.44 - 0.94] | 0.71<br>[0.5 - 1.02] | 0.83<br>[0.44 - 1.22] | 0.69<br>[0.41 - 0.88] | 0.51<br>[0.29 - 0.67] | 0.83<br>[0.47 - 1.23] |
| | $R^2$ | 0.72<br>[-0.64 - 0.88] | 0.74<br>[-1.01 - 0.9] | 0.63<br>[-0.65 - 0.81] | 0.53<br>[-1.24 - 0.78] | 0.81<br>[-0.27 - 0.95] | 0.64<br>[-2.15 - 0.89] |
|  | Max error | 1.63<br>[1.17 - 2.13] | 1.64<br>[1.38 - 2.24] | 1.82<br>[1.18 - 2.69] | 1.62<br>[1.21 - 2.29] | 1.24<br>[0.78 - 1.62] | 1.87<br>[1.03 - 2.68] |
| Lateral<br>Translation<br>[mm] | RMSE | 1.05<br>[0.81 - 1.45] | 0.96<br>[0.77 - 1.45] | 1.23<br>[0.95 - 1.64] | 1.09<br>[0.79 - 1.29] | 1.13<br>[0.58 - 1.8] | 0.81<br>[0.59 - 1.02] |
| | $R^2$ | -0.65<br>[-2.14 - 0.03] | -0.68<br>[-2.84 - 0.07] | -1.47<br>[-6.1 - -0.42] | -1.14<br>[-3.98 - -0.21] | -0.59<br>[-3.35 - -0.14] | -0.06<br>[-1.15 - -0.11] |
|  | Max error | 2.28<br>[1.73 - 2.74] | 2.21<br>[1.67 - 2.6] | 2.6<br>[2.05 - 3.11] | 2.44<br>[2.05 - 2.86] | 2.2<br>[1.33 - 3.56] | 1.99<br>[1.44 - 2.4] |
| Internal<br>Rotation Angle<br>[°] | RMSE | 5.09<br>[3.97 - 6.05] | 5.09<br>[3.45 - 6.17] | 5.43<br>[3.72 - 7.03] | 5.79<br>[4.58 - 7.45] | 2.63<br>[1.58 - 3.73] | 5.97<br>[4.24 - 8.88] |
| | $R^2$ | -1.35<br>[-6.29 - -0.82] | -2.62<br>[-11.02 - -1.12] | -3.12<br>[-15.14 - -1.04] | -3.34<br>[-10.32 - -1.81] | 0.33<br>[-1.09 - 0.32] | -2.41<br>[-10.68 - -3.31] |
|  | Max error | 11.24<br>[8.76 - 12.63] | 11.04<br>[7.74 - 14.31] | 13.13<br>[10.03 - 16.14] | 13.83<br>[10.72 - 16.82] | 5.14<br>[3.47 - 6.6] | 13.76<br>[8.3 - 20.93] |
| Valgus<br>Rotation Angle<br>[°] | RMSE | 1.22<br>[0.99 - 1.68] | 1.25<br>[0.93 - 1.76] | 1.33<br>[0.94 - 1.99] | 1.48<br>[0.94 - 1.99] | 0.75<br>[0.44 - 1.17] | 1.15<br>[0.84 - 1.47] |
| | $R^2$ | -1.71<br>[-5.44 - -0.82] | -1.53<br>[-4.54 - -0.12] | -2.0<br>[-5.03 - -0.6] | -2.76<br>[-9.22 - -2.44] | -0.25<br>[-1.28 - 0.06] | -2.68<br>[-12.09 - -1.67] |
|  | Max error | 2.69<br>[2.35 - 3.27] | 2.79<br>[2.2 - 3.37] | 2.84<br>[2.15 - 3.59] | 3.17<br>[2.03 - 3.95] | 1.75<br>[1.24 - 2.6] | 2.88<br>[1.9 - 3.92] |
| Flexion<br>Rotation Angle<br>[°] | RMSE | 5.01<br>[2.39 - 7.31] | 5.84<br>[2.09 - 9.25] | 5.91<br>[2.18 - 8.73] | 5.0<br>[2.28 - 7.62] | 3.47<br>[1.78 - 4.74] | 3.85<br>[1.62 - 5.65] |
|  | R<sup>2</sup> | 0.97<br>[0.93 - 0.99] | 0.9<br>[0.63 - 0.98] | 0.92<br>[0.8 - 0.99] | 0.97<br>[0.93 - 0.99] | 0.99<br>[0.96 - 1.0] | 0.99<br>[0.97 - 1.0] |
|  | Max error | 7.66<br>[4.78 - 11.41] | 8.32<br>[4.22 - 15.08] | 8.44<br>[5.74 - 12.74] | 7.54<br>[4.74 - 10.81] | 6.43<br>[3.91 - 10.35] | 7.34<br>[3.03 - 11.94] |

### Muscle Activations

During level walking, predicted muscle activations were qualitatively consistent with EMG timing and patterns (Figure 4), except for the rectus femoris.

**Figure 4:**
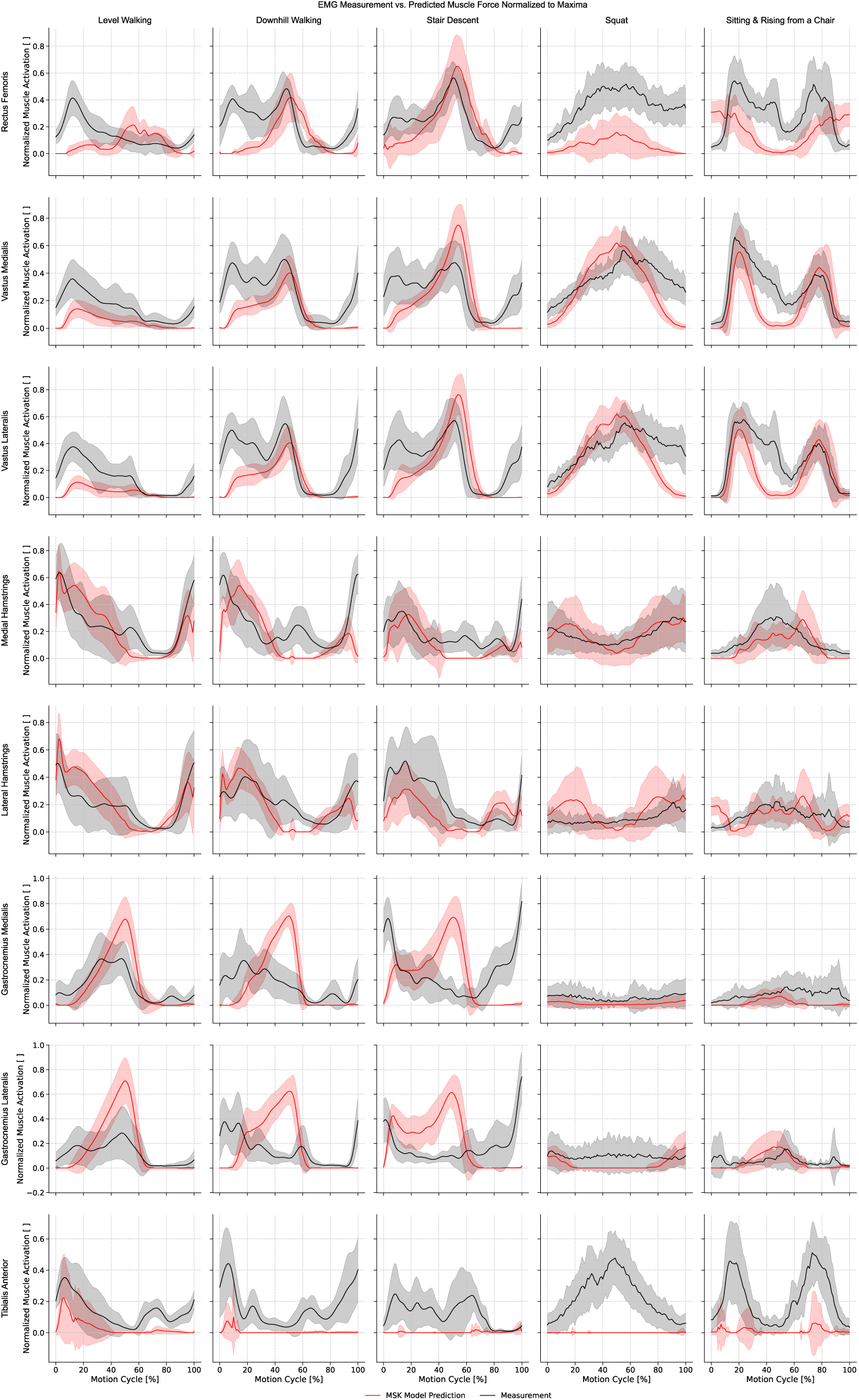
Mean (± SD) predicted muscle forces (red) vs. experimental EMG signals (black) from the CAMS-Knee datasets [32] during different ADLs. Both muscle forces and EMG signals were normalized to the maximum value recorded across all trials and activities for each subject (and limb).

Downhill walking and stair descent presented larger quadriceps EMG signals (vasti and rectus femoris) during the loaded stance phase compared to level walking. The MSK models correctly predicted increased quadriceps activation during push-off but underpredicted it during the swing-to-stance transition and early stance. For the gastrocnemius, the models predicted a large activation during the push-off phase of these activities, which was not observed in the EMG recordings. In contrast, the EMG signals showed a distinct gastrocnemius activation peak in terminal swing that persisted into initial contact, which was not reproduced by the models. The hamstrings showed an activation timing similar to the EMGs, while the tibialis anterior activation was underpredicted.

Squatting, sitting and rising from a chair showed overall good agreement in terms of timing and patterns, except for the tibialis anterior activation which was underpredicted during squatting, and the rectus femoris, which was overpredicted while sitting.

## DISCUSSION

This study established a modelling framework in which MSK model predictions were compared against in vivo measurements from the CAMS Knee dataset, including KCF&Ms, EMG signals, and local knee kinematics. By evaluating model performance across 184 activity cycles from six subjects performing five different ADLs, this work represents the most extensive quantitative validation of an MSK modelling pipeline for predicting knee mechanics to date, expanding upon previous validation efforts that were typically limited to individual activities or smaller trial sets. The proposed modelling approach, incorporating an elastic-foundation contact model between implant components and a surgically balanced ligament model derived from cadaveric testing, reproduced overall knee loading with good agreement and captured qualitative trends in secondary kinematics, while revealing task- and component-specific discrepancies that reflect current limitations in modelling assumptions.

The current results align with previously reported RMSE values for predicted KCFs during level walking and sagittal bending motions. During level walking, our prediction errors for compressive KCFs (i.e., proximo–distal component) (RMSE = 0.29 [0.21 – 0.37] BW) were comparable to the best-performing models based on the *Grand Challenge* datasets that were not directly calibrated to measured KCFs or EMG data [39]. Our results for level-walking are also comparable to the single-subject, single-gait-trial validation reported by Marra et al. [30] (RMSE= 0.26 BW, R^2^ = 0.89), who developed a subject-specific model incorporating CT-derived bone morphology and a FDK implant contact formulation. Esrafilian et al. [63] predicted compressive KCFs during gait variations with an RMSE of approximately 0.43 BW (value digitized from the published graph), despite the use of EMG-assisted MSK models with an elastic foundation contact formulation. Notably, comparable or lower prediction errors were achieved in the present study without EMG-assisted prediction of muscle activations, demonstrating that the proposed modelling framework can reproduce global knee loading during gait without requiring EMG-informed calibration of muscle recruitment. Our predicted muscle activation patterns during level walking were largely consistent with EMG timing, except for the rectus femoris. Surface EMG recordings of this muscle are known to be susceptible to cross-talk from adjacent vasti, and the predicted activation pattern was consistent with previously reported fine-wire EMG measurements [64], suggesting that the discrepancy may reflect EMG measurement limitations rather than modelling inaccuracies.

For squatting, our modelling workflow yielded errors (RMSE = 0.44 [0.20 – 0.88] BW) comparable to those reported by Guitteny et al. [38], who improved squat predictions by constraining the implant contact point trajectories with squat-specific fluoroscopy data, albeit at the expense of reduced accuracy during level walking.

Guo et al. [65] validated their MSK models against data of a single CAMS subject (K5R), for which they had additional access to a full post-operative CT scan of the operated limb, and in which they prescribed the implant flexion-extension angle measured with the moving fluoroscope as input to their simulations. They achieved excellent accuracy in predicted secondary kinematics and reported a RMSE of 0.10 and 0.16 BW for the axial KCF during level walking and squatting, compared to our RMSE of 0.36 and 0.20 BW for the same subject during the same activities. However, prescribing implant motion in one DOF constrains the remaining secondary DOFs, which are intrinsically coupled to the flexion-extension angle due to high conformity of the implant articulating surfaces. In contrast, the present framework was driven exclusively by marker-based motion capture, without prescribing fluoroscopy-derived kinematics, thereby relying solely on motion data typically available in standard motion analysis settings and supporting a more generalizable and broadly applicable modelling approach. As a result, the predicted implant flexion–extension angle reflected the known accuracy limitations of optical motion capture systems [66], resulting in RMSEs of 5.84° and 3.47° during level walking and squatting, respectively. Limitations in kinematic accuracy can propagate to the predicted forces and moments in MSK models [67]. At the same time, the high level of accuracy that Guo et al. [65] obtained in their single subject-study suggests that inclusion of subject-specific bone morphology might be critical to improve MSK models’ performance.

Furthermore, the present analysis evaluated three-dimensional force and moment components, providing a more comprehensive assessment of model performance compared to most previous studies, which, as noted by Guitteny et al. [38], have primarily focused on the proximo–distal compressive force. This broader validation scope revealed that, while global compressive loading was predicted robustly, moderate errors can still be present in other force and moment components.

Regarding implant kinematics, model predictions and fluoroscopic measurements showed qualitatively similar trends across all ADLs, although the level of agreement varied across DOFs. A key strength of this study is the extension of direct quantitative validation beyond level walking and squatting to a wider spectrum of daily activities. Overall, the model predicted the total KCF magnitude with good accuracy across all tasks (RMSE = 0.46 [0.35 – 0.61] BW), although larger errors were observed in certain activities. Stair descent, and, to a lesser extent, downhill walking, exhibited the largest discrepancies during the push-off phase, which were primarily attributable to an overprediction of gastrocnemius activation, highlighting the model’s sensitivity to muscle coordination. Since muscle forces are the primary contributors to knee joint loading [68], deviations in predicted muscle recruitment patterns directly affect the accuracy of the predicted contact forces. Consequently, multi-activity validation not only provides a more stringent assessment of model performance but also serves a diagnostic role by exposing potential issues that would remain undetected if validation were limited to level walking alone. The role of the triceps surae during the push-off phase of stair descent and downhill walking, as well as relevant insights for future improvements in the prediction of these muscles’ activations, are further discussed in Appendix B available in the Supplemental Materials. In particular, modelling gastrocnemius strength as a function of knee flexion substantially reduced the overprediction of gastrocnemius activation during stair descent and led to a normalization of the predicted KCFs, although further work is required to establish a generalized formulation across activities. At the same time, the absence of predicted gastrocnemius activation during the swing phase of stair descent further highlights a known limitation of static optimization approaches, which do not account for anticipatory muscle pre-activation prior to ground contact.

The presented workflow included model personalization based on all subject-specific input data available within the CAMS-Knee datasets. Importantly, the modelling framework was implemented without activity- or subject-specific tuning or calibration to match the experimental measurements, such that all results reflect a single, consistently applied set of modelling assumptions across tasks and participants. Nevertheless, several sources of modelling uncertainty remain, including (1) the lack of subject-specific bone morphology and muscle lines of action, which were instead represented using the refined TLEM 2 cadaveric template, (2) the positioning of the implant components in certain DOFs, which was assumed from implant-specific surgical guidelines, and (3) the behaviour of the ligamentous structures surrounding the knee, which was derived from in-house cadaveric data. These uncertainties likely contribute to the observed inter-subject variability in the accuracy of our models’ predictions. Due to the lack of intra-operative subject-specific ligament information, the ligament model was derived from a single cadaveric specimen, thereby avoiding the introduction of arbitrary subject-specific ligament modelling assumptions into the presented results. The contribution of the soft tissue model (JEF) to KCF&Ms was qualitatively small (Appendix C available in the Supplemental Materials), indicating that ligaments are minor contributors compared to muscles, external loads, and contact between implant components. This observation may be specific to the INNEX ultra- congruent design investigated here and could differ in less constrained or cruciate-retaining implants, where preserved ligament structures may play a more prominent role in joint mechanics. The uncertainty associated with ligament modelling warrants further investigation through dedicated sensitivity analyses. Recent developments in anatomically based ligament modelling, including the identification of ligament attachment sites and near-isometric fibers from joint kinematics [69], provide an alternative approach that could be explored in future studies.

Another limitation of this study is that the patellar motion was modelled as a function of the knee flexion angle, rather than as a fully articulated joint. Future work should therefore focus on subject-specific, FDK-based modelling of the patellofemoral joint, as this can influence tibiofemoral mechanics, particularly at deep flexion angles [38, 70]. It should also be noted that the FDK approach does not account for friction between implant components [54], and that its robustness across different implant designs remains to be evaluated.

While model personalization remains advisable [30, 65, 71], implementing it all at once may conceal limitations in generic modelling assumptions. In the present study, incremental, stepwise personalization enabled the identification of critical features requiring further refinement, such as the line of action of the Achilles and the patellar tendons, and an incorrect activation pattern of the triceps surae during stair descent and downhill walking (Appendix A and B available in the Supplemental Materials). In this sense, the presented validation framework can serve as a meaningful benchmark to support generic model improvements and staged personalization.

Although this study documents the development of a validation tool to assess the accuracy of MSK models, it does not determine the minimum level of accuracy that is needed for translation into specific clinical applications. Such information remains to be determined, for instance by establishing quantifiable correlations between patient outcomes and biomechanical variables of interest. Recent literature suggests that local knee kinematics, knee moments, and muscle coordination during different ADLs might be best suited for identifying poor post-operative functionality and clinical outcomes [72–74].

## CONCLUSION

While previous MSK modelling studies have mostly focused on validating predictions during level walking and occasionally during squatting, the current study investigated a broader range of ADLs and subjects and therefore represents the most extensive MSK quantitative validation for the prediction of knee loading and local kinematics to date. The development and continuous refinement of accurate MSK models will enable us to better understand how knee mechanics can influence patient satisfaction and mobility during real-life activities, while paving the way for dynamic, movement-informed, patient-specific surgical planning.

## APPENDIX

More information on the study is provided in the Supplemental Materials.

## ACKNOWLEDGMENT

The authors would like to thank Eik Siggelkow (Zimmer Biomet) for his valuable support in the implementation of the cadaveric-derived ligament model and the elastic-foundation contact model.

