## Supplementary Material for "Extending Musculoskeletal Modelling Validation Beyond Gait: Knee Loading and Kinematics During Daily Activities"

**Appendix A: TLEM 2.2 geometrical refinement**

In the early stages of this work, simulations based on the generic 2.1 lower-limb model led to the identification of critical musculoskeletal geometrical features with incorrect anatomical representation. This appendix documents the working logic and simulation techniques for identifying specific model features in need of refinement and provides a description of the implemented modifications. The implemented refinements have been incorporated in the TLEM 2.2 model, which has been released as part of the AnyBody Model Managed Repository (AMMR, https://anyscript.org/ammr/changelog.html) and served as a base for the simulations reported in the main article.

The MSK models based on the TLEM 2.1 [1] lower-limb model were scaled to match anthropometrics and CT-landmark data of the CAMS subjects, similar to what is described in the main article. Virtual implantation was also performed to define the instrumented-tibial-tray-based coordinate system ($CS_{JWI\_TIB}$, [2]), in which KCFs were reported. The knee joint however was modelled as the default 1-DOF hinge, with the knee flexion axis defined using a functional approach derived from kinematic recordings obtained during passive, unloaded range-of-motion testing of the cadaveric specimen in the TLEM 2.0 dataset [3].

*Muscle contributions to joint moments and forces*

Muscle path uncertainties arising from estimated insertion points, wrapping geometries, and inaccurate local scaling can lead to errors in predicted joint loads [1]. Possible critical geometries were identified by analysing the contributions of individual muscle elements to the three vector components of the KCFs and to the net joint moments [1, 4, 5]. The contribution of each joint-spanning muscle to the KCF components was computed by projecting the muscle force vectors onto $CS_{JWI\_TIB}$ and normalized by BW.

Muscle contributions to the net joint moments in each anatomical plane were computed as the product of the force exerted by the muscle times the distance of its instantaneous line of action from the centre of rotation of the joint. Muscle contribution to the net joint moments were computed in the default joint coordinate systems in AMS, which are aligned with joint axes of rotation thus providing a more functional interpretation of muscle function, and normalized by body mass.

Muscles that presented a high contribution to a KCF component in a direction perpendicular to the main anatomical plane they spanned, or muscles that provided a joint moment contribution larger than the overall joint net moment, or a joint moment contribution antagonist to the net joint moment during a given motion, were flagged as muscles that could potentially present errors in their geometrical definition and were further inspected.

*Achilles tendon distal wrapping*

In the TLEM 2.1 gastrocnemii (medialis and lateralis) and soleus did not share the same distal line of action, with the gastrocnemii presenting a more favourable lever arm around the ankle, particularly in dorsiflexed positions (Fig. S1). This led the gastrocnemii to provide a relatively larger plantarflexing contribution to the ankle plantarflexion moment compared to soleus, particularly during the push-off phase of stair descent (Fig. S2). At the same time the gastrocnemii provided an antagonist flexing contribution to the knee extension moment, which in turn had to be counteracted by a larger extension contribution by the vasti, causing high compressive forces at the knee. A more realistic wrapping of the distal lines of action of gastrocnemius medialis and lateralis, soleus, and plantaris was introduced around ankle and calcaneus. Wrapping cylinders ensured that these muscles ran parallel along the distal section of the tibia, resembling the Achilles tendon anatomy, and provided equal contributions to the ankle plantarflexion moment (Fig. S3), thus partially reducing the overprediction in KCFs during stair descent. Pre- and post-refinement muscle plantarflexion lever arms are reported (Fig. S4).

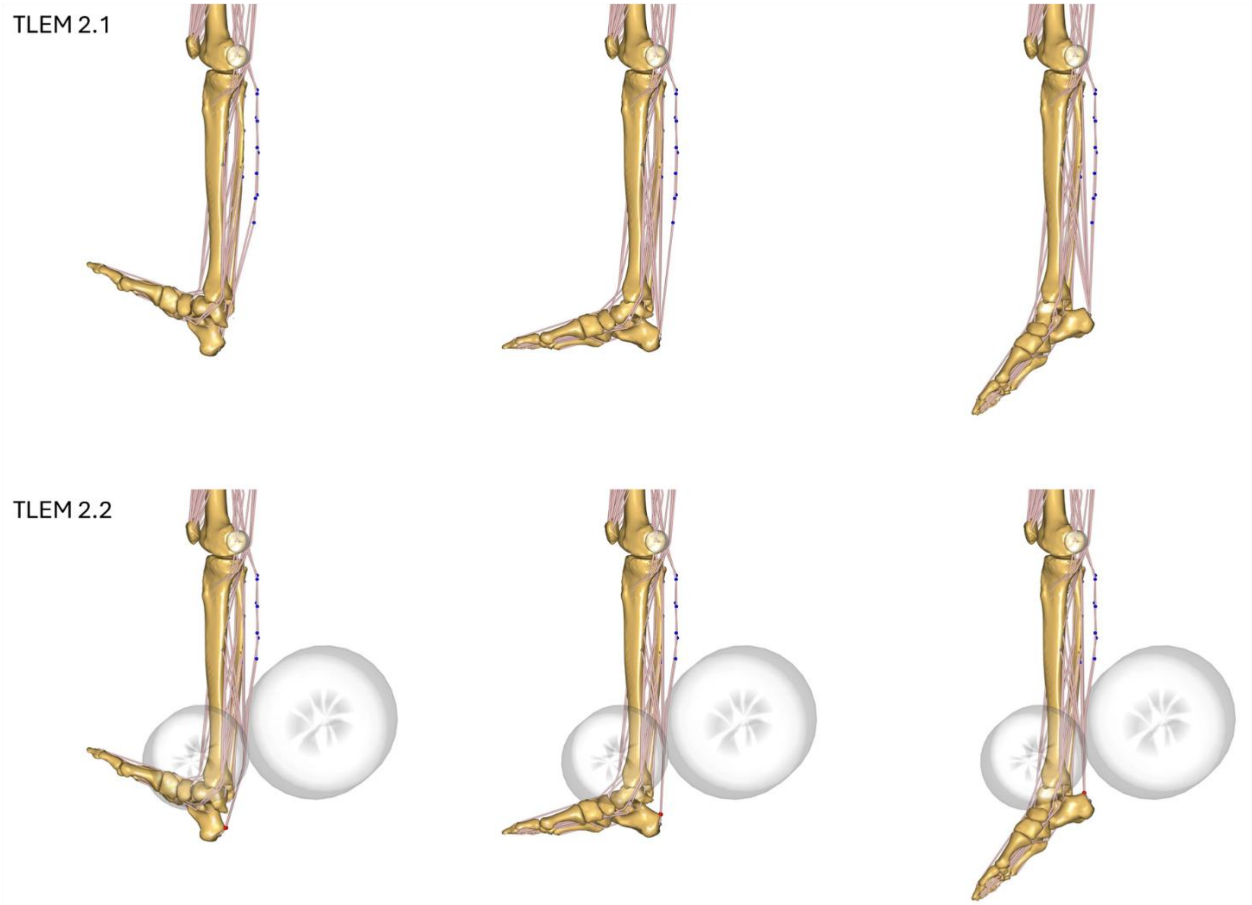


*Figure S1: Lines of action of gastrocnemius and soleus at different ankle flexion angles with associated via points (in blue) and wrapping cylinders.*

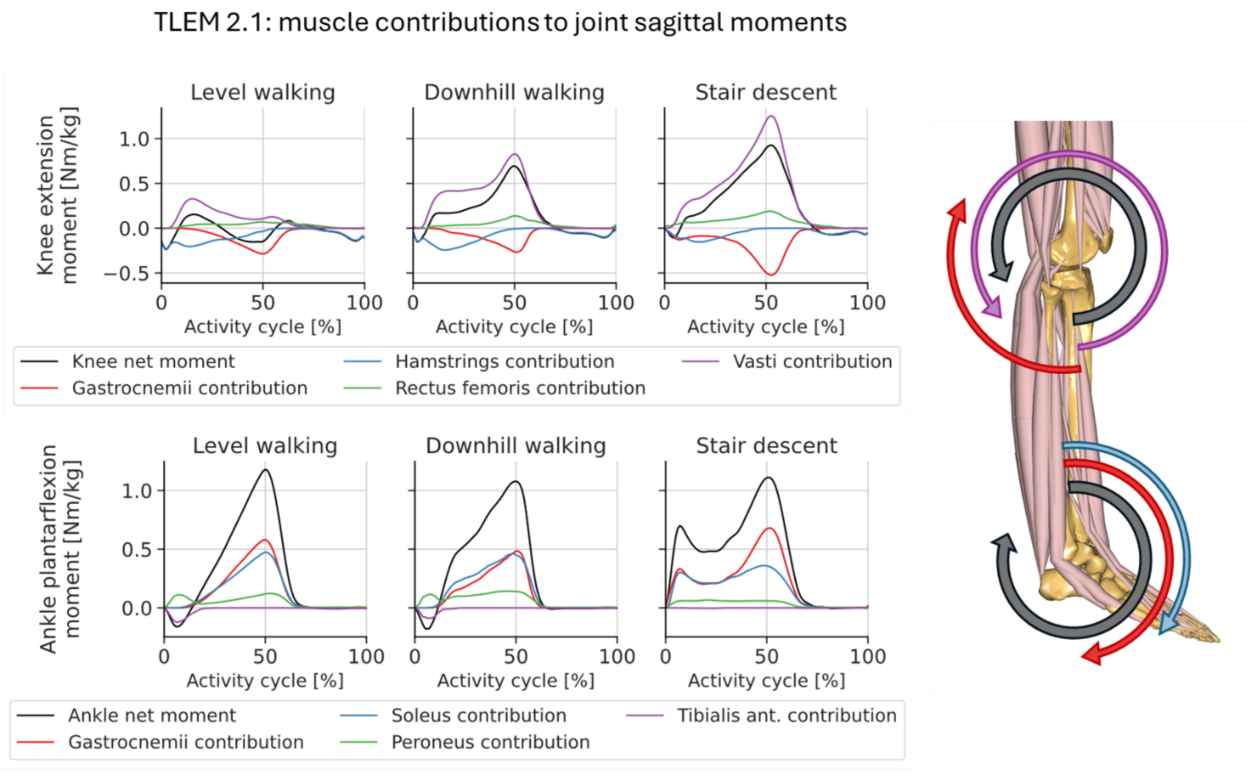


*Figure S2: Muscle contributions to knee and ankle sagittal moments for the TLEM 2.1 model. Mean value across all subjects and trials.*

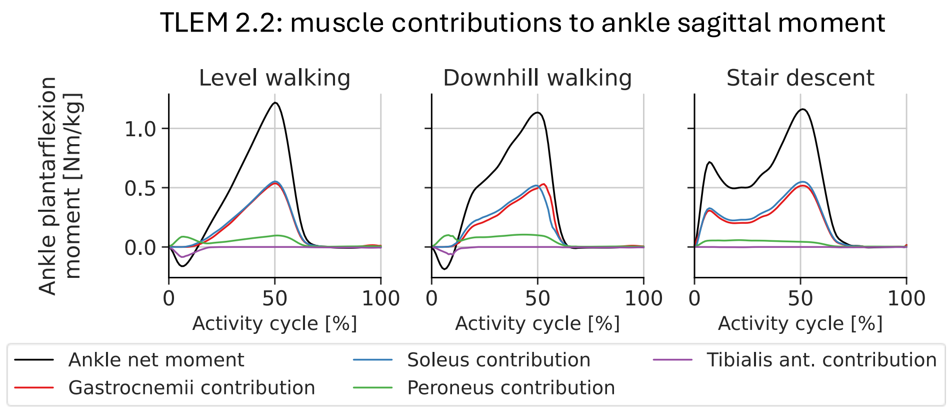


*Figure S3: Muscle contributions to the ankle sagittal moment for the TLEM 2.2 model. Mean value across all subjects and trials.*

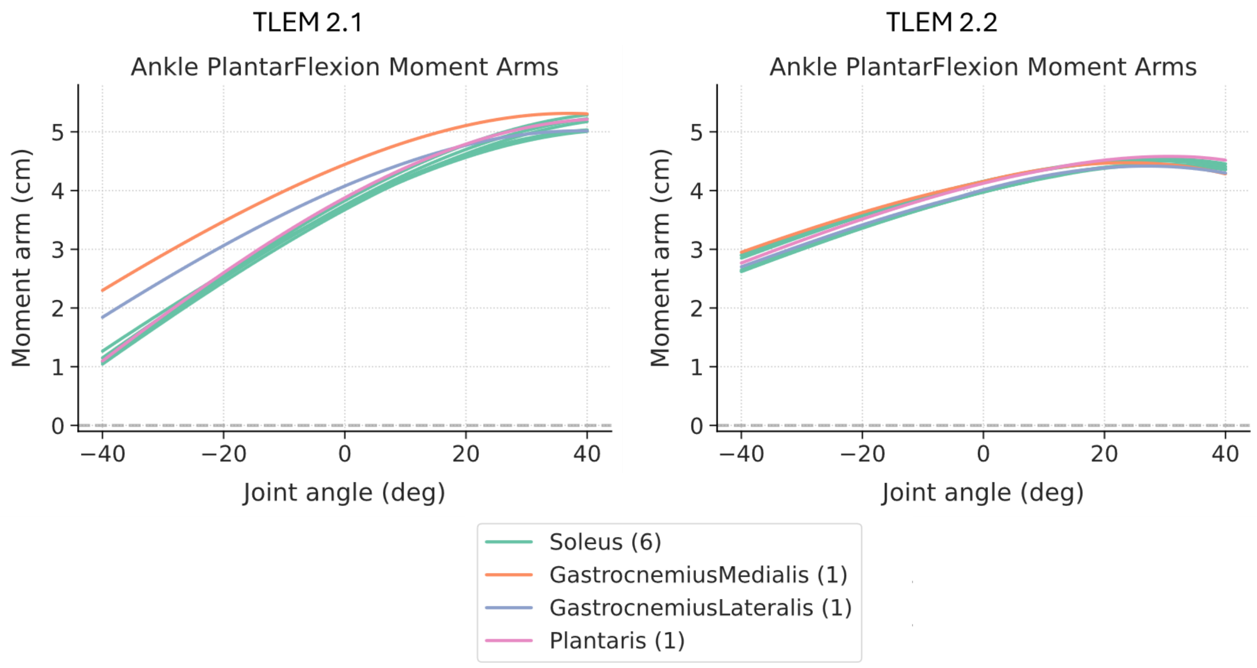


*Figure S4: Ankle plantar-flexors' sagittal moment arms over a range of ankle flexion for the TLEM 2.1 and 2.2.*

*Transversal plane knee flexion axis alignment and patellar tendon orientation*

The muscle contribution study revealed a large contribution of the vasti to the lateral component of KCFs across all activities (Fig. S5). While the vasti act mainly as knee flexors in the sagittal plane, the tilted line of action of the patellar tendon in the TLEM 2.0/2.1 geometry (Fig. S6) transmitted their force as a laterally oriented shear force at the knee level. The introduction of a 20° internal rotation offset in the definition of the knee axis in the proximal tibia compared to the cadaveric-measured axis (from the TLEM 2.0) led to a neutral alignment between femur and tibia in the transversal plane and to a longitudinally oriented line of action of the patellar tendon throughout the range of motion of the knee.

While KCF data from implanted knees may not be considered the most adequate reference data to evaluate the anatomical alignment of the native tibio-femoral joint complex, we just interpreted the discrepancy in the predicted KCFs as a flag to further investigate the definition of the native anatomical joint against anatomical atlases. The model was not tuned nor overfitted to match the CAMS-Knee experimental data.

The knee flexion axis of the original TLEM 2.0 dataset was defined using a functional approach derived from kinematic recordings obtained during passive, unloaded range-of-motion testing of the cadaveric specimen. The flexion axis was then registered relative to bony landmarks from a straight-leg CT scan of the cadaveric specimen. The relative position between femur and tibia during the CT scan was maintained as the neutral extension pose in early iterations of the model. We suspect that, during the CT scan, the tibia was externally rotated relative to the femur, due to either the absence of muscle tension or passive joint conformity (i.e., the screw home mechanism). This likely introduced a rotational alignment offset in the neutral extension pose. This rotational offset persisted throughout the range of motion, as the passive, unloaded cadaveric testing failed to capture the natural external-to-internal rotation of the tibia at deeper flexion angles. The implemented modification should therefore better describe the behaviour of the knee joint throughout the range of knee flexion, with a more realistic longitudinal orientation of the patellar tendon that does not lead to excessive laterally oriented shear forces.

We recommend future users to always verify the modelling assumptions regarding the knee joint, as they could affect the outcomes of their specific use-cases. Whenever possible, a personalization of the knee axis of rotation remains advisable [6-10], as was performed in the main article with the implementation of an elastic foundation contact model based on implant geometry.

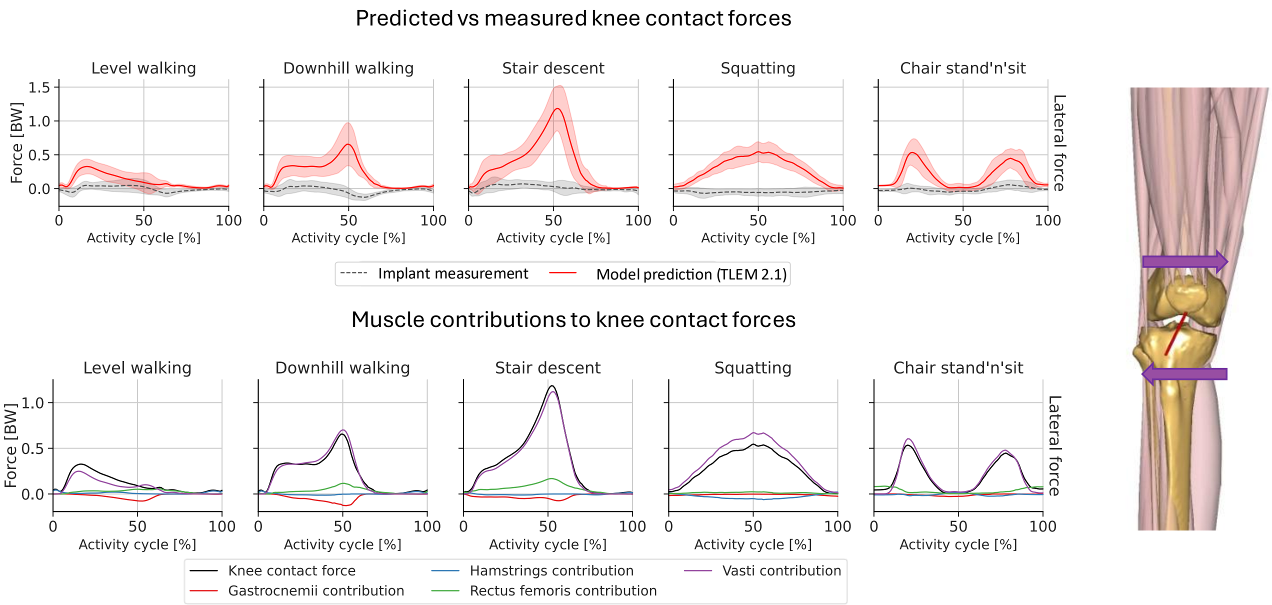


*Figure S5: Predicted vs measured lateral knee contact forces, and relative contributions of different muscles to the predicted values.*

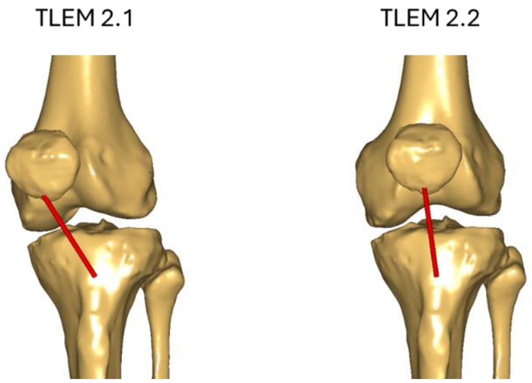


*Figure S6: Patellar tendon orientation in the TLEM 2.1 and 2.2.*

**Appendix B: Triceps surae activation during downhill walking and stair descent**

Despite the geometrical refinements described above and the implementation of an advanced FDK knee joint based on implant geometry, an overprediction in KCF&Ms in the sagittal plane (proximal and anterior KCFs and extension KCM) during the push-off phase of downhill walking and stair descent persisted. This overprediction corresponds with an excessive gastrocnemii activation compared to the EMG data.

The experimental EMG data from the CAMS-Knee datasets shows a distinct activation for the gastrocnemii between level walking, where the muscles activate in midstance and push-off, and stair descent, where they activate just before heel strike but are minimally activated during midstance and push-off. This activation pattern is consistent with previously reported EMG data for healthy subjects [11-14]. Yet, the overall kinetic requirements at the ankle are similar between level walking, downhill walking, and stair descent, with all the activities requiring the generation of a comparable net ankle plantarflexion moment during push-off (Fig. S3, black line).

From the EMG data we know that the gastrocnemii should not activate during the push-off phase of stair descent. Furthermore, given the biarticular nature of the gastrocnemii, any active or passive force that is generated by the gastrocnemii muscle-tendon units will lead to an increase in KCFs, which would not be consistent with the experimental data from the instrumented implants. At the same time, soleus EMG data from literature [12-14] suggest that this muscle also presents limited activation during the push-off phase of stair descent. This leaves the passive components of the soleus muscle-tendon units, or other uniarticular passive structures at the ankle, as the most plausible providers of the required plantarflexion moment during stair descent push-off, when the ankle is in a highly dorsiflexed position.

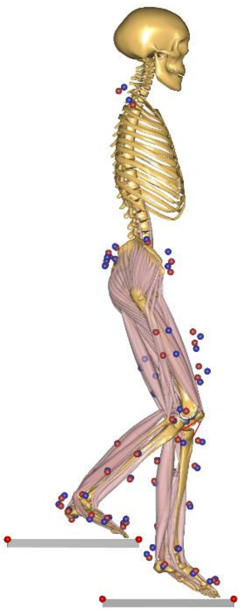

*Figure S7: Push off phase during stair descent with a dorsiflexed ankle (trailing leg).*

In addition to the models' predictions described in the main article – which are here referred to as "baseline" – four additional triceps surae models were tested and compared: A) gastrocnemii and soleus were modelled as Hill-type actuators, whose optimal muscle fiber and tendon slack lengths were calibrated with default routines in AMS; B) the calibration poses were adapted so that the soleus presents maximal strength in a dorsiflexed position and the gastrocnemii in a plantarflexed position, thus making the soleus more favourably recruited during stair descent push-off; C) a recruitable ankle torque actuator, which conceptually represents the uniarticular soft tissue structures capable of passively resisting ankle dorsiflexion (soleus bundles of the Achilles tendon, posterior joint capsule and ligaments), was implemented in parallel to constant-strength triceps surae muscles. The actuator could be recruited to provide plantarflexion torque when the ankle is dorsiflexed more than 20°; D) the strength of the gastrocnemii was modelled as knee-flexion dependent regardless of the ankle position, starting to decrease above 20° of knee flexion and became null above 30°, while the soleus strength remained constant.

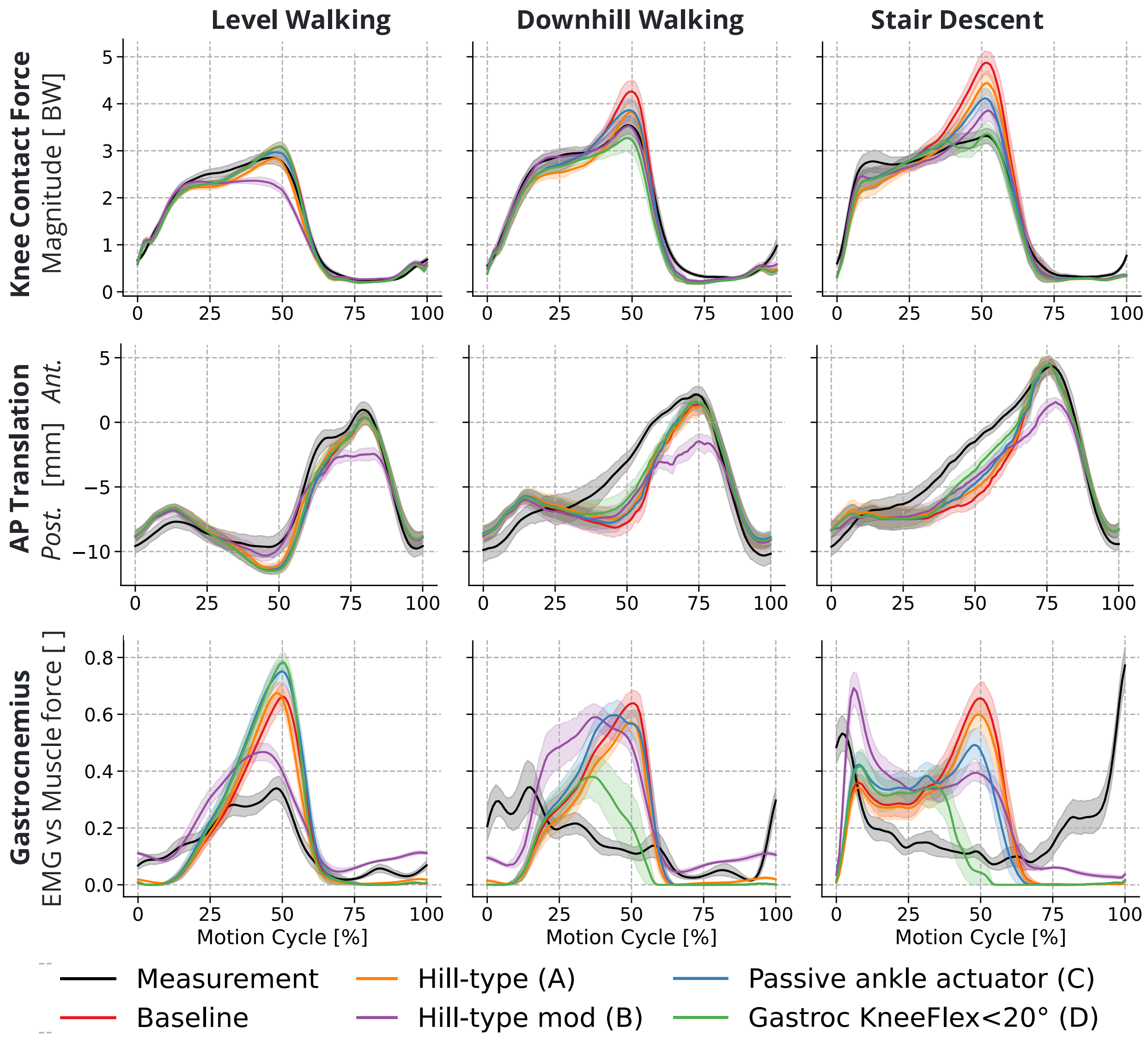


*Figure S8: Mean (± 95 CI) KCF magnitude, anterior posterior translation of the femoral component with respect to the tibial baseplate, and gastrocnemius activation during different ADLs for different triceps surae models.*

Modelling the triceps surae as Hill-type actuators (Model A, Fig. S8 in orange) only lead to minor differences compared to the baseline results. Adapting the calibration poses of the gastrocnemii and soleus (Model B, in violet) partially reduced the activation of the gastrocnemii during stair descent push-off, but at the same time led to a reduction in gastrocnemii activation during level walking, resulting in an underprediction of KCFs and AP translations. The implementation of an ankle torque actuator (Model C, in blue) only led to a small reduction in peak gastrocnemii activation. This indicates that modelling passive anatomical structures may not be sufficient to predict the correct activation of the triceps surae. When modelling the gastrocnemii strength as knee-flexion-dependent (Model D, in green) we could correctly predict a reduced gastrocnemii activation during stair-descent push-off, leading to a normalization of the predicted KCFs. Despite representing an approximation of the real behaviour of the gastrocnemii, this solution suggests that understanding the recruitment of the gastrocnemii themselves is crucial to model stair descent correctly. Lauber et al. [15] proposed that the relationship between gastrocnemius motor units' activation and knee flexion could be attributed to variations in muscle fiber length and could therefore be explainable with a Hill-type muscle model. Herzog et al. [16] also suggested that the gastrocnemii operate in the ascending part of the typical Hill-type strength profile *in vivo*. However, further analysis of gastrocnemius muscle tendon units (MTU) length and contraction/elongation velocity during different ADLs did not show any clear difference between activities (Fig. S9), suggesting that a Hill-type muscle model might not be sufficient to discriminate different gastrocnemii activation patterns during different ADLs. While our models solved the muscle redundancy problem through a 3$^{rd}$-oder-polynomial recruitment criterion, other central nervous system recruitment strategies might be at play here. For instance, Suzuki et al. [17] showed that, during an isometric ankle plantarflexion task on a dynamometer, the addition of a voluntary isometric knee extension moment depressed the activation of the knee-extension-antagonist gastrocnemius medialis, despite its length remaining constant throughout the isometric task, and favoured the activation of the soleus instead. This inhibitory behaviour cannot be predicted with the cost functions that are typically used to solve the muscle redundancy problem in inverse-dynamics-based MSK modelling workflows, which aim at minimizing the sum of squared or cubed muscle activations [18]. Additional terms in the cost function might be required to model muscle co-activation around specific joints and/or the activity modulation of biarticular muscles like the gastrocnemii [19-21].

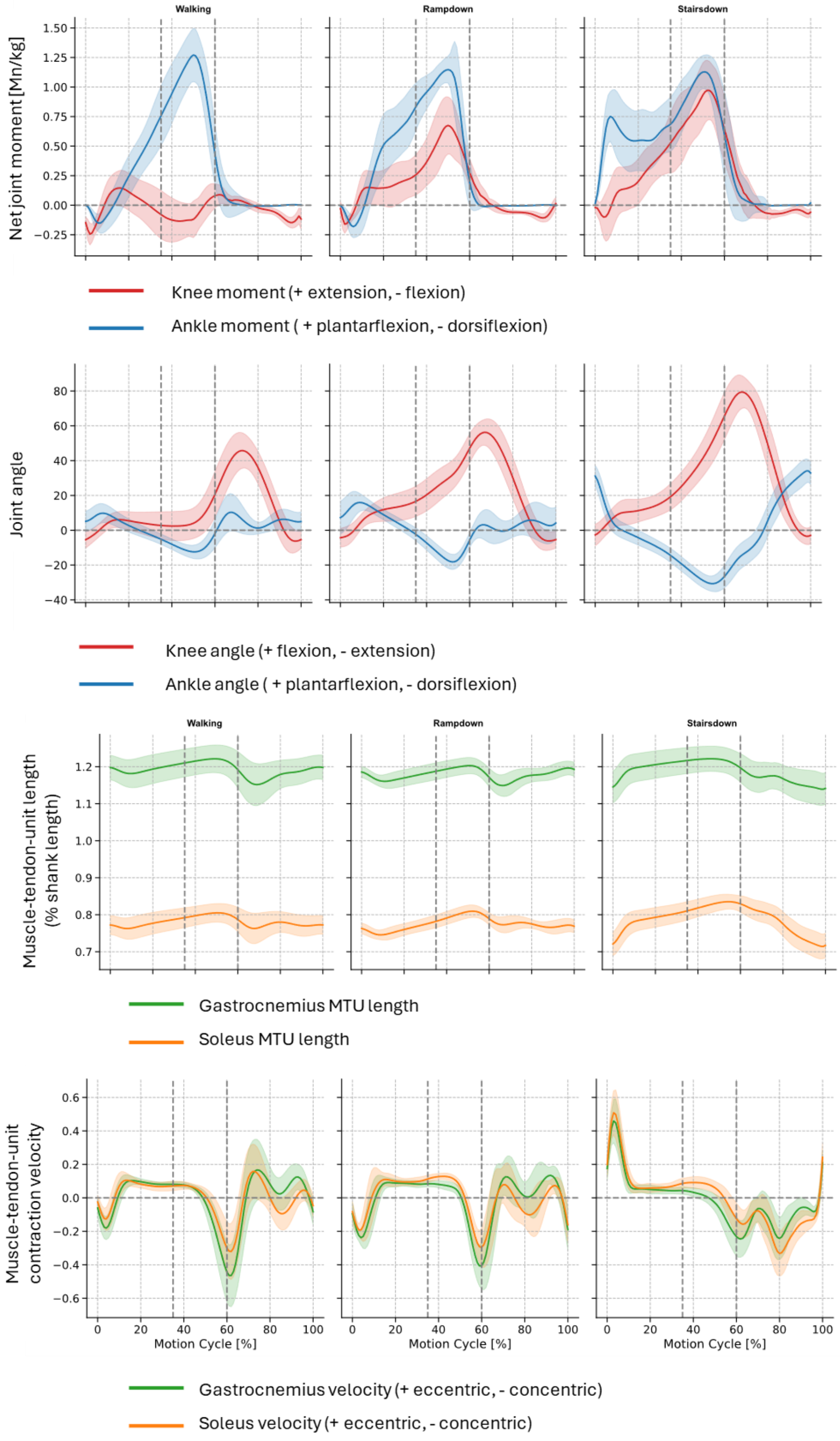


*Figure S9: knee and ankle net joint moments and angles during level walking, downhill walking and stair descent. Soleus and gastrocnemius muscle tendon unit (MTU) length is normalized as a percentage of the shank segment length. MTU velocity is the first derivative of the length and indicates whether the muscle is elongating (+) or shortening (-).*

Stair descent was seldomly analyzed with MSK modelling in the literature so far [12, 22]. Valente et al. [22] observed an overprediction of the gastrocnemii similar to the one discussed in this study, while Lin et al. [12] correctly predicted a low activation of the gastrocnemii during stair descent push-off, but reported an overprediction of the soleus activation. Current models lack the capability to accurately predict the activation of the triceps surae in a consistent way across different activities of daily living, particularly during stair descent. Modifications aiming to reduce the gastrocnemius strength when the knee is flexed, such as the one presented here, can lead to improvements in KCFs prediction during stair descent and downhill walking without leading to any further discrepancy during other ADLs (Figs. S10, S11, S12). While this approach could be sufficiently accurate for certain applications looking at overall knee joint loading, the resulting models are still far from accurately predicting the force generated by the individual muscles and their relative substructures, both active and passive.

A generalized triceps surae model that consistently works across different activities and tasks would most certainly require a detailed characterization of the muscle tendon architecture and the tissue properties (e.g., compliance of the Achilles tendon bundles) [23], as well as a better understanding of the central nervous system recruitment strategy for these specific muscles.

In the absence of a generalized triceps surae model, EMG-informed simulations might bypass our limited understanding of gastrocnemii and soleus recruitment and mitigate the currently observed issues [24-26]. However, Esrafilian et al. [26] predicted compressive KCF during variations of level walking with a RMSE of approximatively 0.43*BW (estimated from graph digitization) despite using of EMG-assisted MSK models with an elastic foundation contact formulation, suggesting that EMG-assistance alone might not be sufficient to drastically improve predictions. Furthermore, the calibration of the EMG-assisted muscle properties led to a decrease in maximum strength of the gastrocnemius and an increase for the soleus [26], consistently with what is suggested here for a more conventional (static optimization) approach to solve the muscle recruitment redundancy problem. Previous static-optimization-based analyses have also suggested the introduction of different weighing factors for gastrocnemii and soleus, as well as for other biarticular and uniarticular muscles, to modulate KCF predictions [26-28]. Similarly, Marra et al. [10] resorted to a 35% reduction in peak muscle strength for all knee extensors and flexors, including the gastrocnemii, thus altering the competitive recruitment of soleus and gastrocnemii to plantarflex the ankle during their analysis of level walking. These solutions, however, are still far from providing us with a better understanding of the central nervous system recruitment strategy for these specific muscles. Predictive simulations of human movement might shed some light on this topic [20, 21, 29].

To conclude, modelling level walking alone is not sufficient to test the boundaries of validity of current MSK models. Future MSK modelling studies should aim at reproducing the in-vivo behaviour of the triceps surae during various activities of daily living as well as during controlled dynamometric tasks [15, 17, 30]. Publicly available motion capture datasets that include stair negotiation [31-33] represent a great opportunity in this regard.

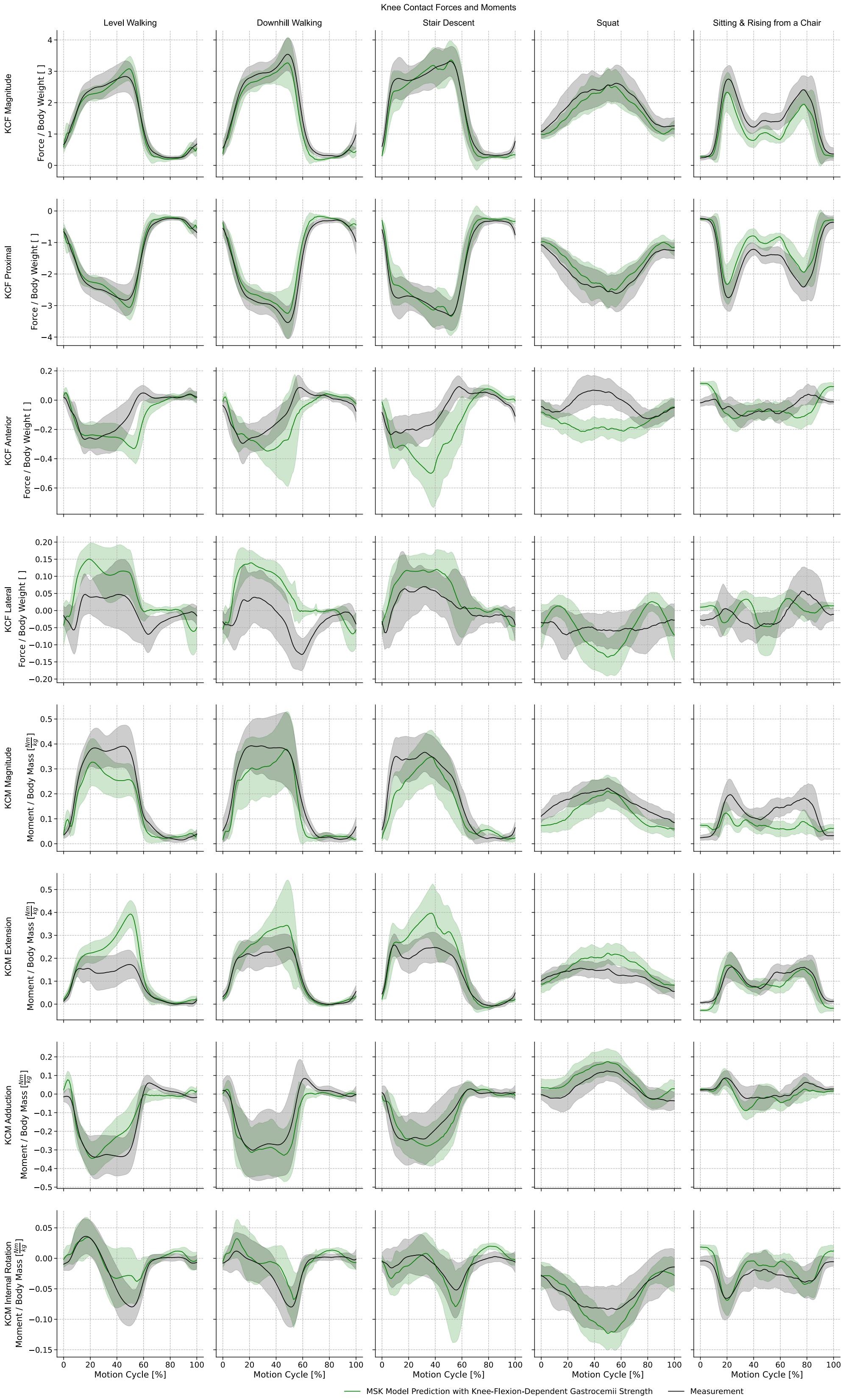


*Figure S10: Mean (± SD) predicted knee contact forces and moments (KCF&Ms; red) when modelling the gastrocnemii strength as knee-flexion-dependent (Gastroc KneeFlex<20°), compared with experimental measurements (black) from the CAMS-Knee datasets during different ADLs.*

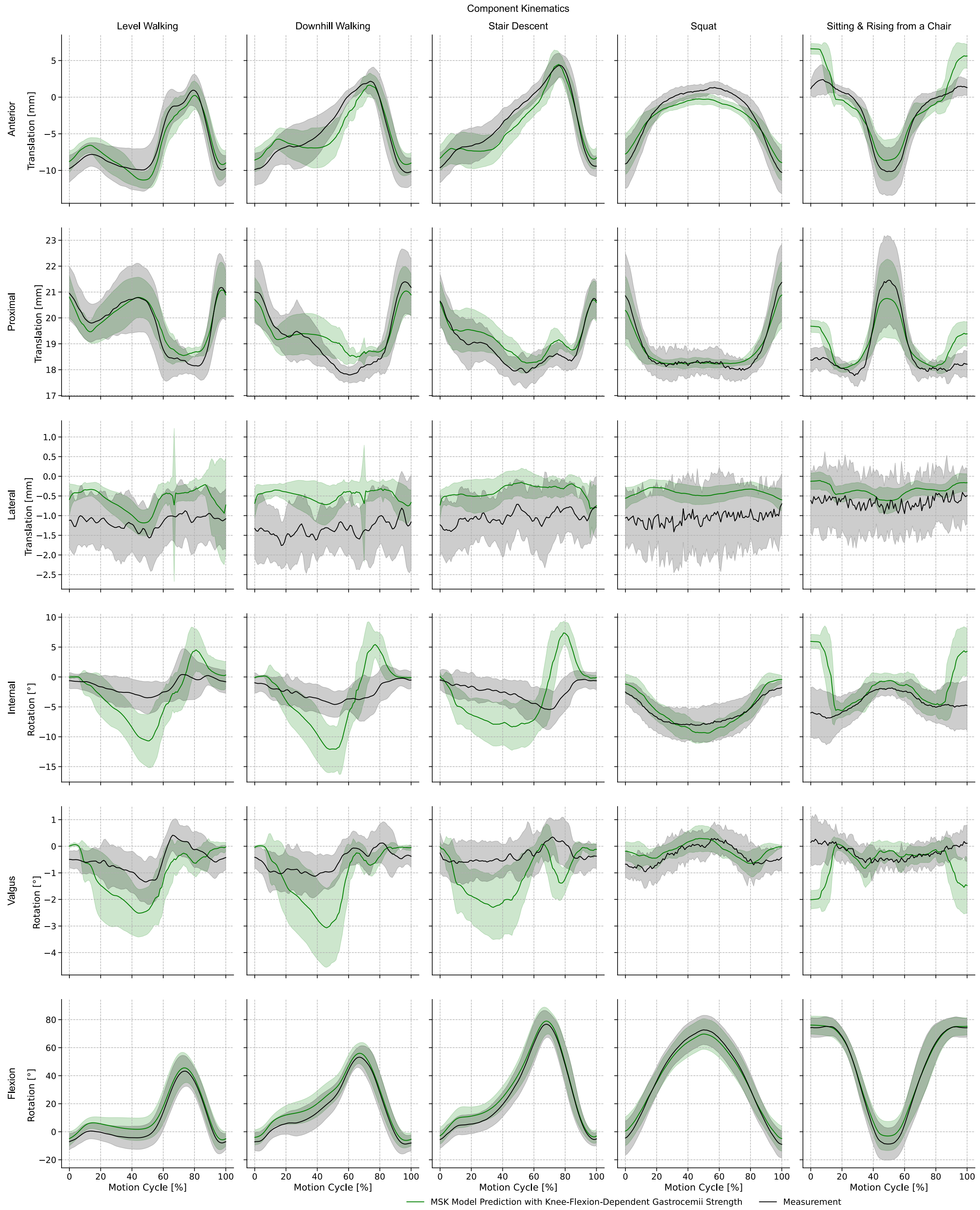


*Figure S11: Mean (± SD) predicted implant kinematics (red) when modelling the gastrocnemii strength as knee-flexion-dependent (Gastroc KneeFlex<20°), compared with experimental measurements (black) from the CAMS-Knee datasets during different ADLs.*

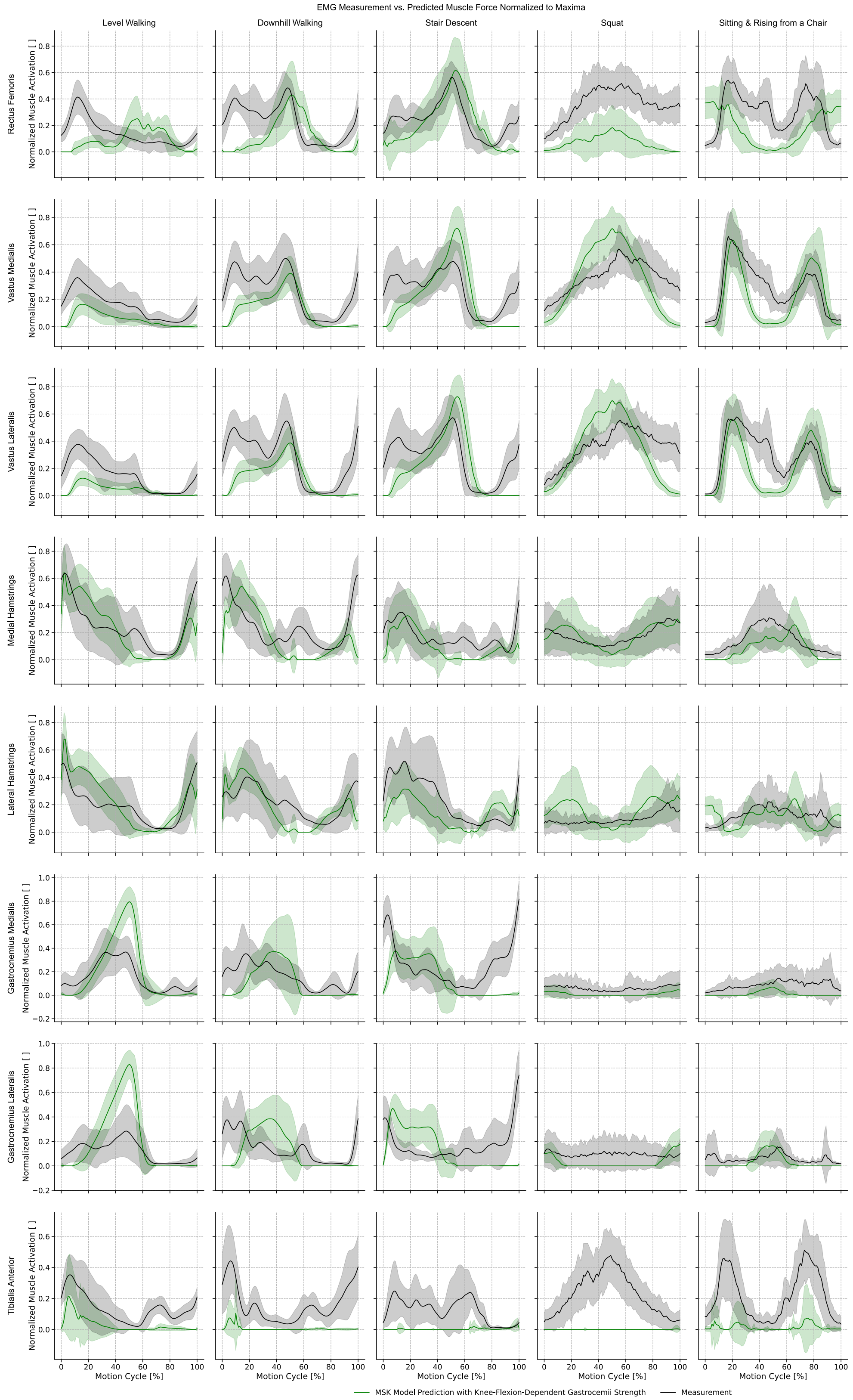


*Figure S12: Mean (± SD) predicted muscle forces (red) when modelling the gastrocnemii strength as knee-flexion-dependent (Gastroc KneeFlex<20°), compared with experimental EMG signals (black) from the CAMS-Knee datasets during different ADLs.*

[4] De Pieri, E., Romkes, J., Wyss, C., Brunner, R., and Viehweger, E., 2022, "Altered Muscle Contributions are Required to Support the Stance Limb During Voluntary Toe-Walking," Frontiers in Bioengineering and Biotechnology, 10.

[5] De Pieri, E., Cip, J., Brunner, R., Weidensteiner, C., and Alexander, N., 2023, "The functional role of hip muscles during gait in patients with increased femoral anteversion," Gait & Posture, 100, pp. 179-187.

[6] De Pieri, E., Nüesch, C., Pagenstert, G., Viehweger, E., Egloff, C., and Mündermann, A., 2022, "High tibial osteotomy effectively redistributes compressive knee loads during walking," Journal of Orthopaedic Research, n/a(n/a).

[14] McFadyen, B. J., and Winter, D. A., 1988, "An integrated biomechanical analysis of normal stair ascent and descent," Journal of Biomechanics, 21(9), pp. 733-744.
[15] Lauber, B., Lichtwark, G. A., and Cresswell, A. G., 2014, "Reciprocal activation of gastrocnemius and soleus motor units is associated with fascicle length change during knee flexion," Physiol Rep, 2(6).
[16] Herzog, W., Read, L. J., and ter Keurs, H. E. D. J., 1991, "Experimental determination of force-length relations of intact human gastrocnemius muscles," Clinical Biomechanics, 6(4), pp. 230-238.
[17] Suzuki, T., Chino, K., and Fukashiro, S., 2014, "Gastrocnemius and soleus are selectively activated when adding knee extensor activity to plantar flexion," Human Movement Science, 36, pp. 35-45.
[18] Rasmussen, J., Damsgaard, M., and Voigt, M., 2001, "Muscle recruitment by the min/max criterion — a comparative numerical study," Journal of Biomechanics, 34(3), pp. 409-415.
[19] Erdemir, A., McLean, S., Herzog, W., and van den Bogert, A. J., 2007, "Model-based estimation of muscle forces exerted during movements," Clinical Biomechanics, 22(2), pp. 131-154.
[20] De Groote, F., and Falisse, A., 2021, "Perspective on musculoskeletal modelling and predictive simulations of human movement to assess the neuromechanics of gait," Proceedings of the Royal Society B: Biological Sciences, 288(1946), p. 20202432.
[21] Dembia, C. L., Bianco, N. A., Falisse, A., Hicks, J. L., and Delp, S. L., 2021, "OpenSim Moco: Musculoskeletal optimal control," PLOS Computational Biology, 16(12), p. e1008493.
[22] Valente, G., Grenno, G., Dal Fabbro, G., Zaffagnini, S., and Taddei, F., 2023, "Medial and lateral knee contact forces during walking, stair ascent and stair descent are more affected by contact locations than tibiofemoral alignment in knee osteoarthritis patients with varus malalignment," Frontiers in Bioengineering and Biotechnology, Volume 11 - 2023.
[23] Finni, T., and Vanwanseele, B., 2023, "Towards modern understanding of the Achilles tendon properties in human movement research," Journal of Biomechanics, 152, p. 111583.
[24] Pizzolato, C., Lloyd, D. G., Sartori, M., Ceseracciu, E., Besier, T. F., Fregly, B. J., and Reggiani, M., 2015, "CEINMS: A toolbox to investigate the influence of different neural control solutions on the prediction of muscle excitation and joint moments during dynamic motor tasks," Journal of Biomechanics, 48(14), pp. 3929-3936.
[25] Meyer, A. J., Patten, C., and Fregly, B. J., 2017, "Lower extremity EMG-driven modeling of walking with automated adjustment of musculoskeletal geometry," PLOS ONE, 12(7), p. e0179698.
[26] Esrafilian, A., Smith, C., Lavikainen, J., Stenroth, L., Mononen, M. E., Karjalainen, P. A., Saxby, D. J., Lloyd, D. G., and Korhonen, R. K., 2025, "An Electromyography-Assisted Musculoskeletal Simulation With Concurrent Optimization of Muscle Excitations and Knee Joint Kinematics," Journal of Biomechanical Engineering, 147(9).
[27] DeMers, M. S., Pal, S., and Delp, S. L., 2014, "Changes in tibiofemoral forces due to variations in muscle activity during walking," Journal of Orthopaedic Research, 32(6), pp. 769-776.
[28] Smith, C. R., Brandon, S. C. E., and Thelen, D. G., 2019, "Can altered neuromuscular coordination restore soft tissue loading patterns in anterior cruciate ligament and menisci deficient knees during walking?," Journal of Biomechanics, 82, pp. 124-133.
[29] Falisse, A., Serrancolí, G., Dembia, C. L., Gillis, J., Jonkers, I., and De Groote, F., 2019, "Rapid predictive simulations with complex musculoskeletal models suggest that diverse healthy

and pathological human gaits can emerge from similar control strategies," Journal of The Royal Society Interface, 16(157), p. 20190402.
[30] Whittington, B., Silder, A., Heiderscheit, B., and Thelen, D. G., 2008, "The contribution of passive-elastic mechanisms to lower extremity joint kinetics during human walking," Gait & Posture, 27(4), pp. 628-634.
[31] Komnik, I., David, S., Funken, J., Haberer, C., Potthast, W., and Weiss, S., 2018, "Compromised knee internal rotation in total knee arthroplasty patients during stair climbing," PLOS ONE, 13(10), p. e0205492.
[32] Lencioni, T., Carpinella, I., Rabuffetti, M., Marzegan, A., and Ferrarin, M., 2019, "Human kinematic, kinetic and EMG data during different walking and stair ascending and descending tasks," Scientific Data, 6(1), p. 309.
[33] Taylor, W. R., Schütz, P., Bergmann, G., List, R., Postolka, B., Hitz, M., Dymke, J., Damm, P., Duda, G., Gerber, H., Schwachmeyer, V., Hosseini Nasab, S. H., Trepczynski, A., and Kutzner, I., 2017, "A comprehensive assessment of the musculoskeletal system: The CAMS-Knee data set," Journal of Biomechanics, 65, pp. 32-39.

**Appendix C: Results for individual patients**

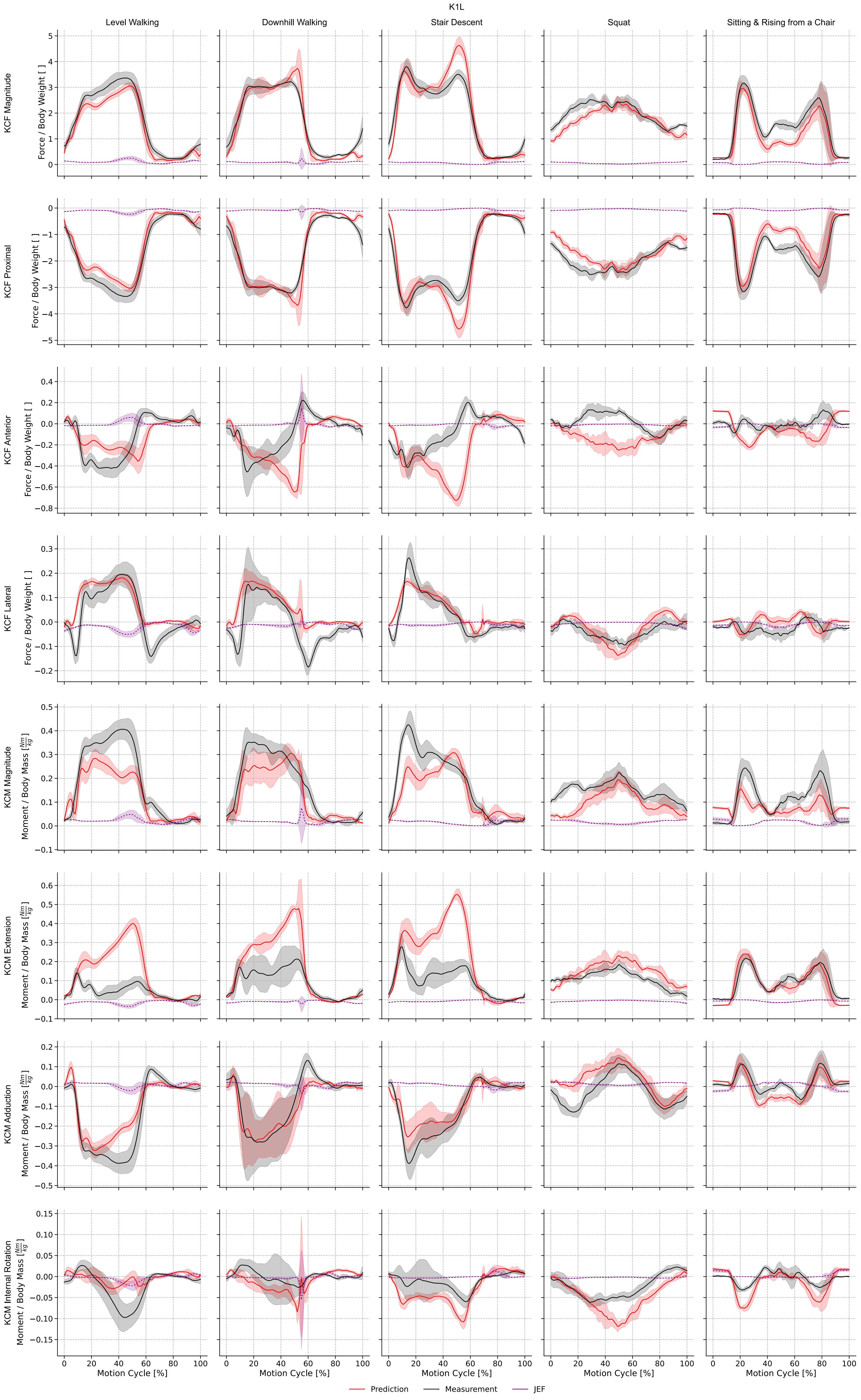

K1L
Level Walking
Downhill Walking
Stair Descent
Squat
Sitting & Rising from a Chair
KCF Magnitude
Force / Body Weight [ ]
KCF Proximal
Force / Body Weight [ ]
KCF Anterior
Force / Body Weight [ ]
KCF Lateral
Force / Body Weight [ ]
KCM Magnitude
Moment / Body Mass [Nm/kg]
KCM Extension
Moment / Body Mass [Nm/kg]
KCM Adduction
Moment / Body Mass [Nm/kg]
KCM Internal Rotation
Moment / Body Mass [Nm/kg]
Motion Cycle [%]
Prediction
Measurement
JEF

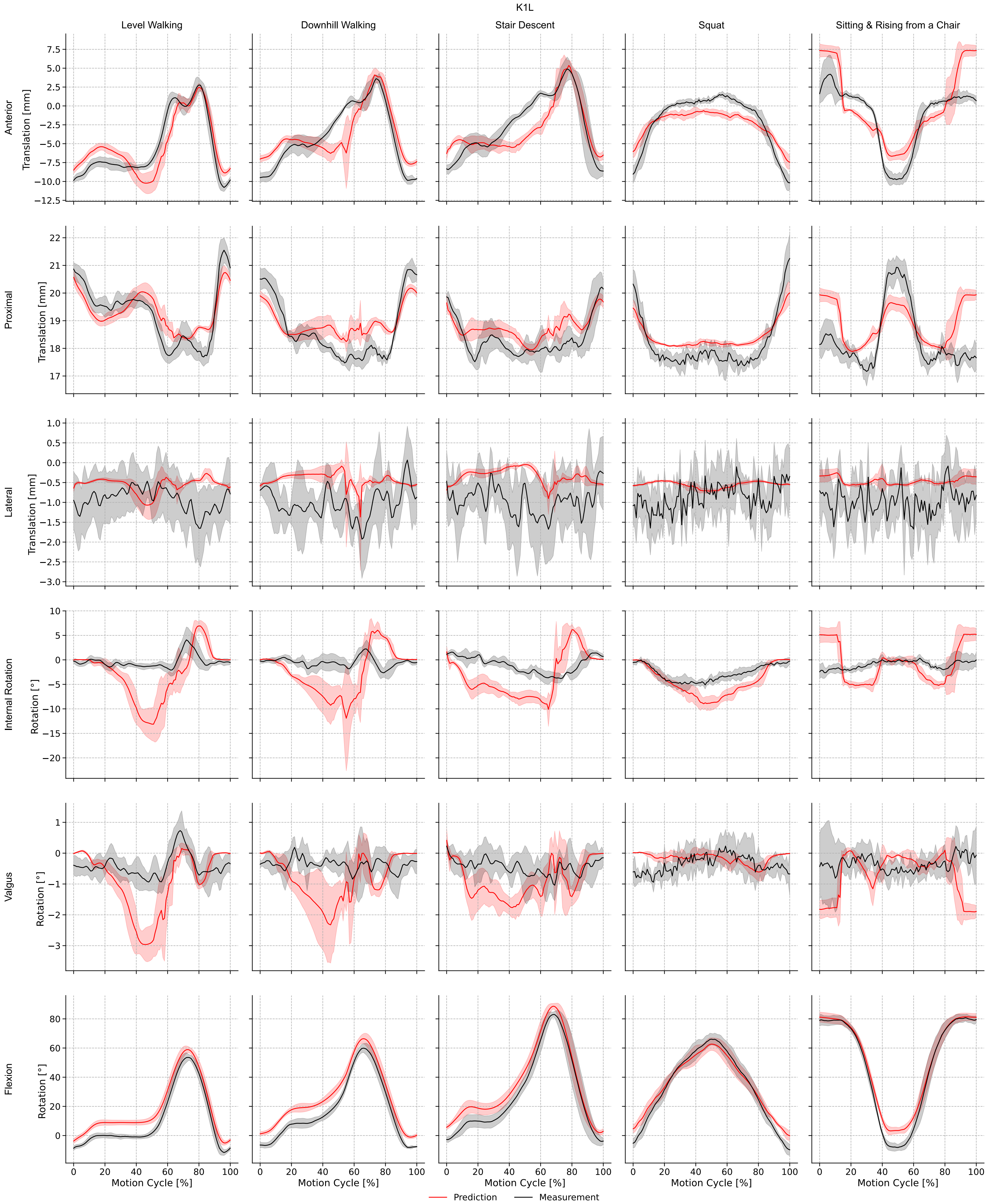

K1L
Level Walking
Downhill Walking
Stair Descent
Squat
Sitting & Rising from a Chair
Anterior
Translation [mm]
Proximal
Translation [mm]
Lateral
Translation [mm]
Internal Rotation
Rotation [°]
Valgus
Rotation [°]
Flexion
Rotation [°]
Motion Cycle [%]
Prediction
Measurement

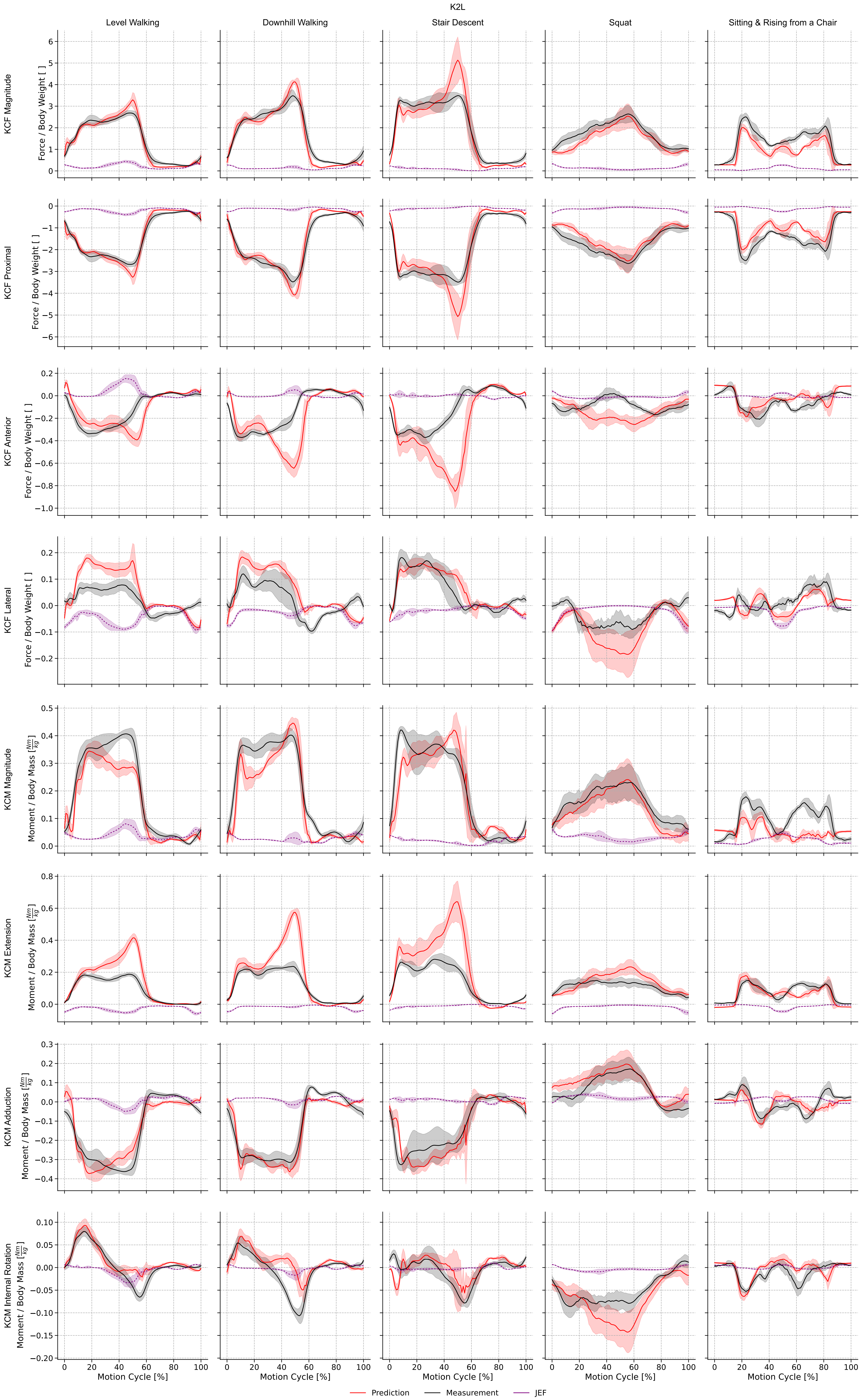
K2L
Level Walking
Downhill Walking
Stair Descent
Squat
Sitting & Rising from a Chair
KCF Magnitude
Force / Body Weight [ ]
KCF Proximal
KCF Anterior
KCF Lateral
KCM Magnitude
Moment / Body Mass [Nm/kg]
KCM Extension
KCM Adduction
KCM Internal Rotation
Motion Cycle [%]
Prediction
Measurement
JEF

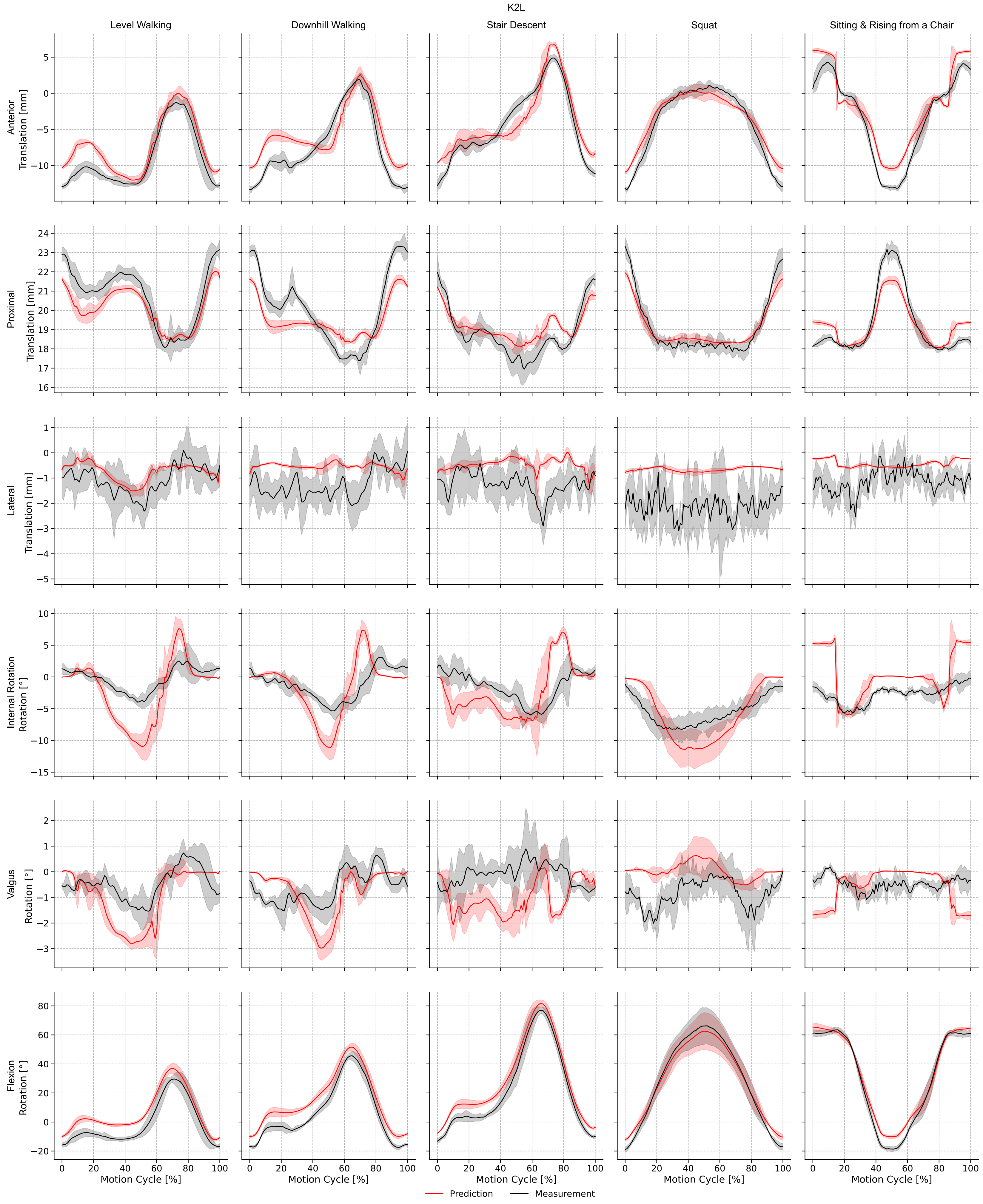
K2L
Level Walking
Downhill Walking
Stair Descent
Squat
Sitting & Rising from a Chair
Anterior Translation [mm]
Proximal Translation [mm]
Lateral Translation [mm]
Internal Rotation Rotation [°]
Valgus Rotation [°]
Flexion Rotation [°]
Motion Cycle [%]
Prediction
Measurement

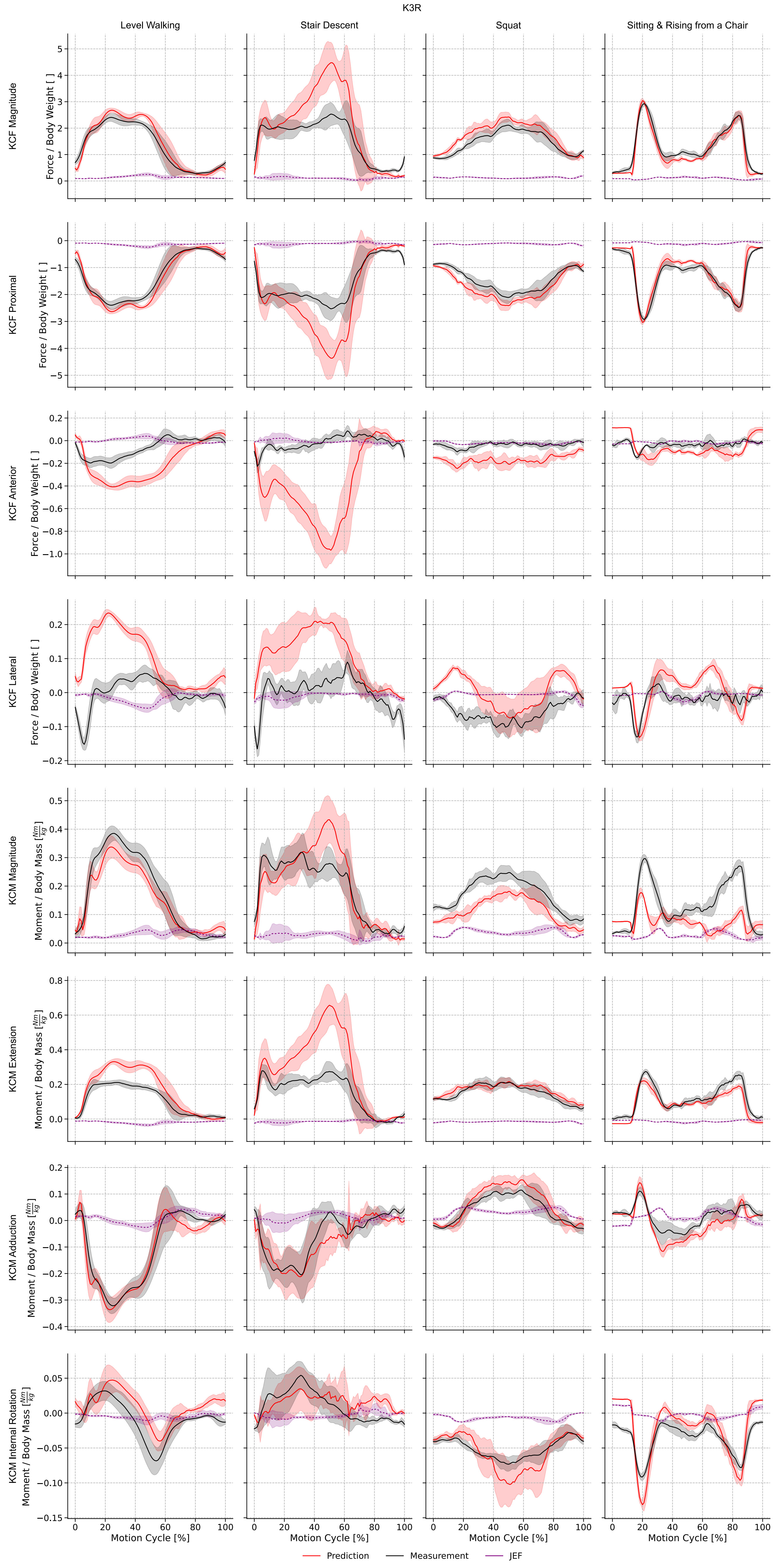
K3R
Level Walking
Stair Descent
Squat
Sitting & Rising from a Chair
KCF Magnitude
Force / Body Weight [ ]
KCF Proximal
Force / Body Weight [ ]
KCF Anterior
Force / Body Weight [ ]
KCF Lateral
Force / Body Weight [ ]
KCM Magnitude
Moment / Body Mass [Nm/kg]
KCM Extension
Moment / Body Mass [Nm/kg]
KCM Adduction
Moment / Body Mass [Nm/kg]
KCM Internal Rotation
Moment / Body Mass [Nm/kg]
Motion Cycle [%]
Prediction
Measurement
JEF

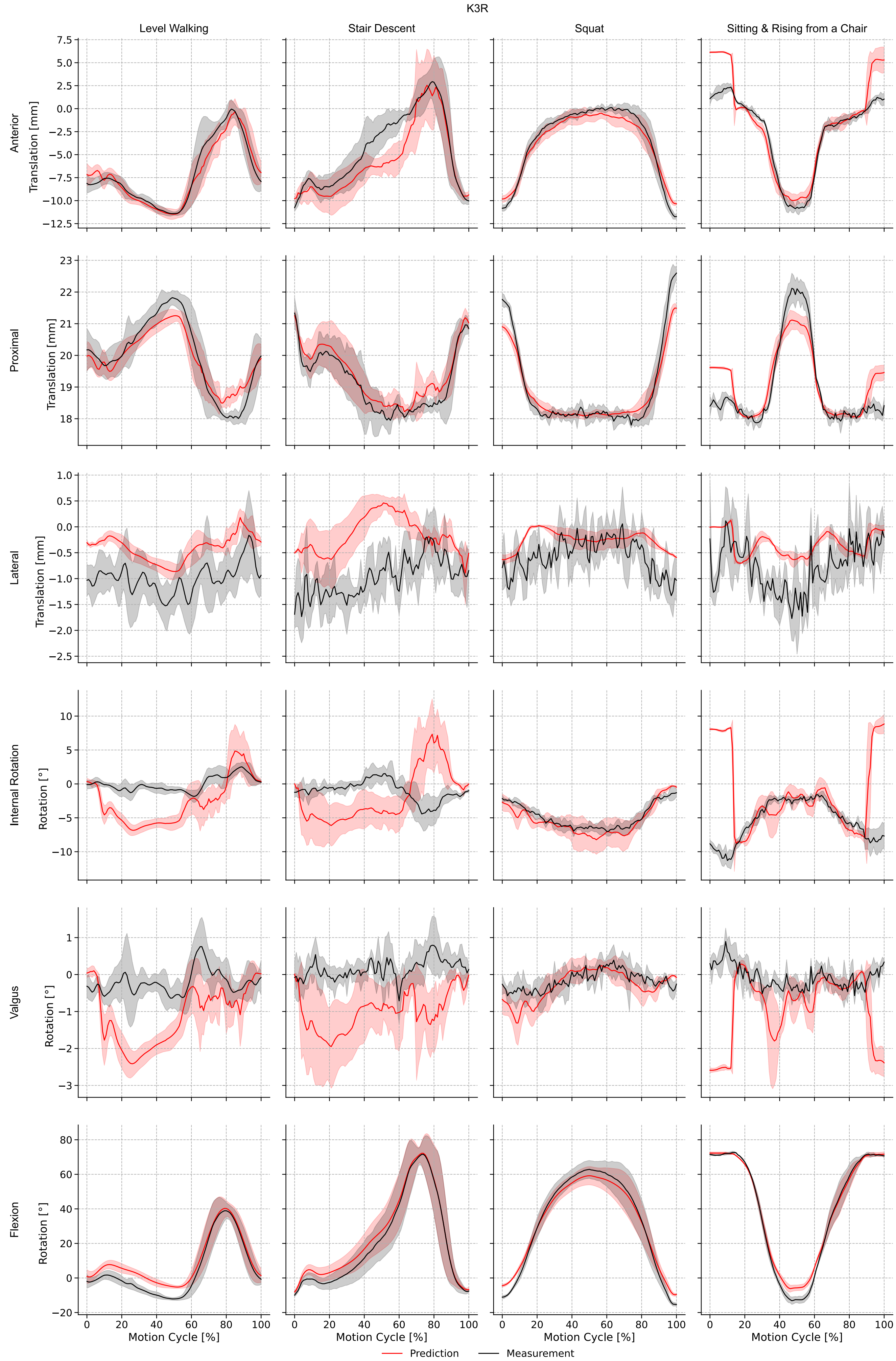
K3R
Level Walking
Stair Descent
Squat
Sitting & Rising from a Chair
Anterior
Translation [mm]
Proximal
Translation [mm]
Lateral
Translation [mm]
Internal Rotation
Rotation [°]
Valgus
Rotation [°]
Flexion
Rotation [°]
Motion Cycle [%]
Prediction
Measurement

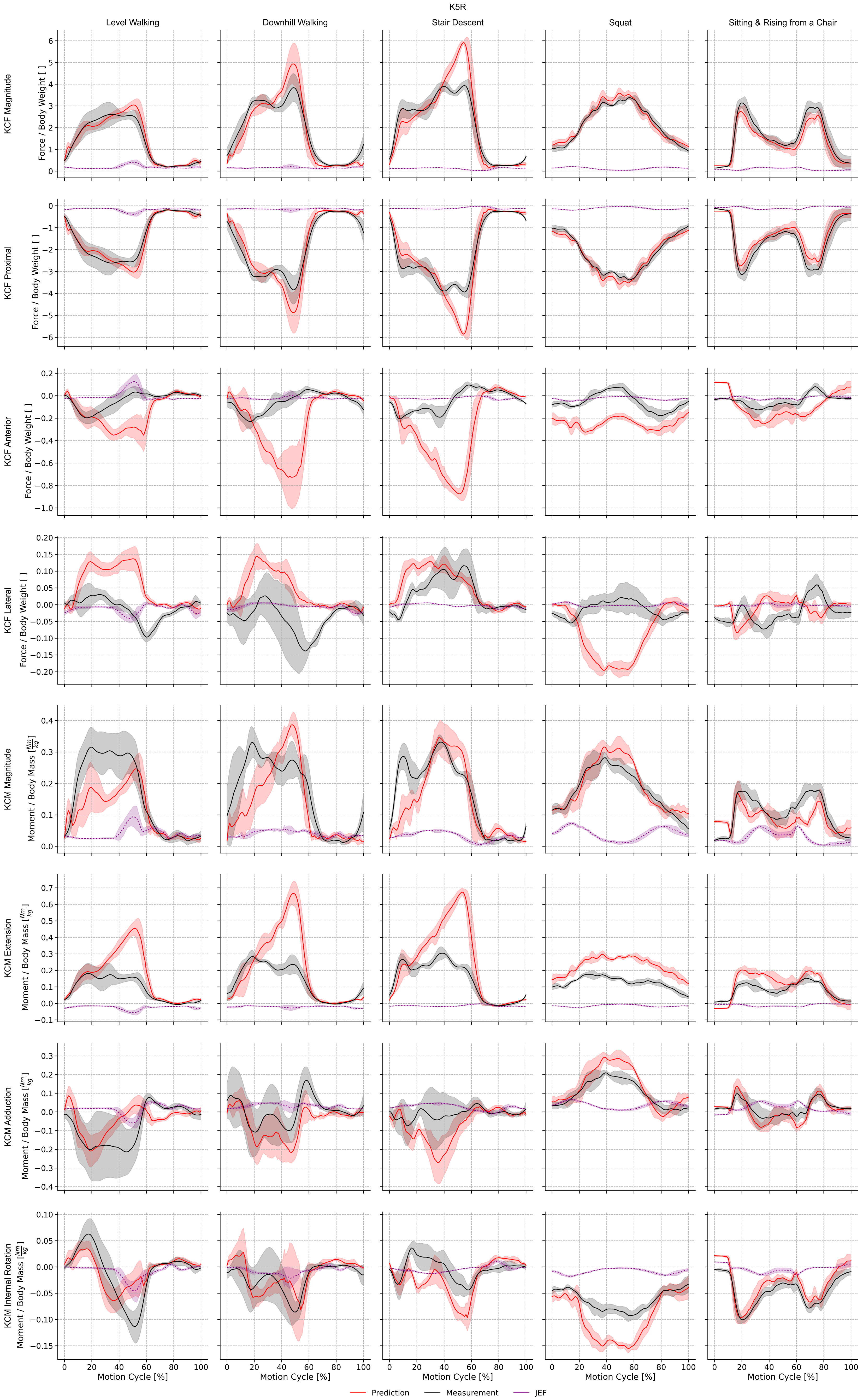

K5R
Level Walking
Downhill Walking
Stair Descent
Squat
Sitting & Rising from a Chair
KCF Magnitude
Force / Body Weight [ ]
KCF Proximal
Force / Body Weight [ ]
KCF Anterior
Force / Body Weight [ ]
KCF Lateral
Force / Body Weight [ ]
KCM Magnitude
Moment / Body Mass [Nm/kg]
KCM Extension
Moment / Body Mass [Nm/kg]
KCM Adduction
Moment / Body Mass [Nm/kg]
KCM Internal Rotation
Moment / Body Mass [Nm/kg]
Motion Cycle [%]
Prediction
Measurement
JEF

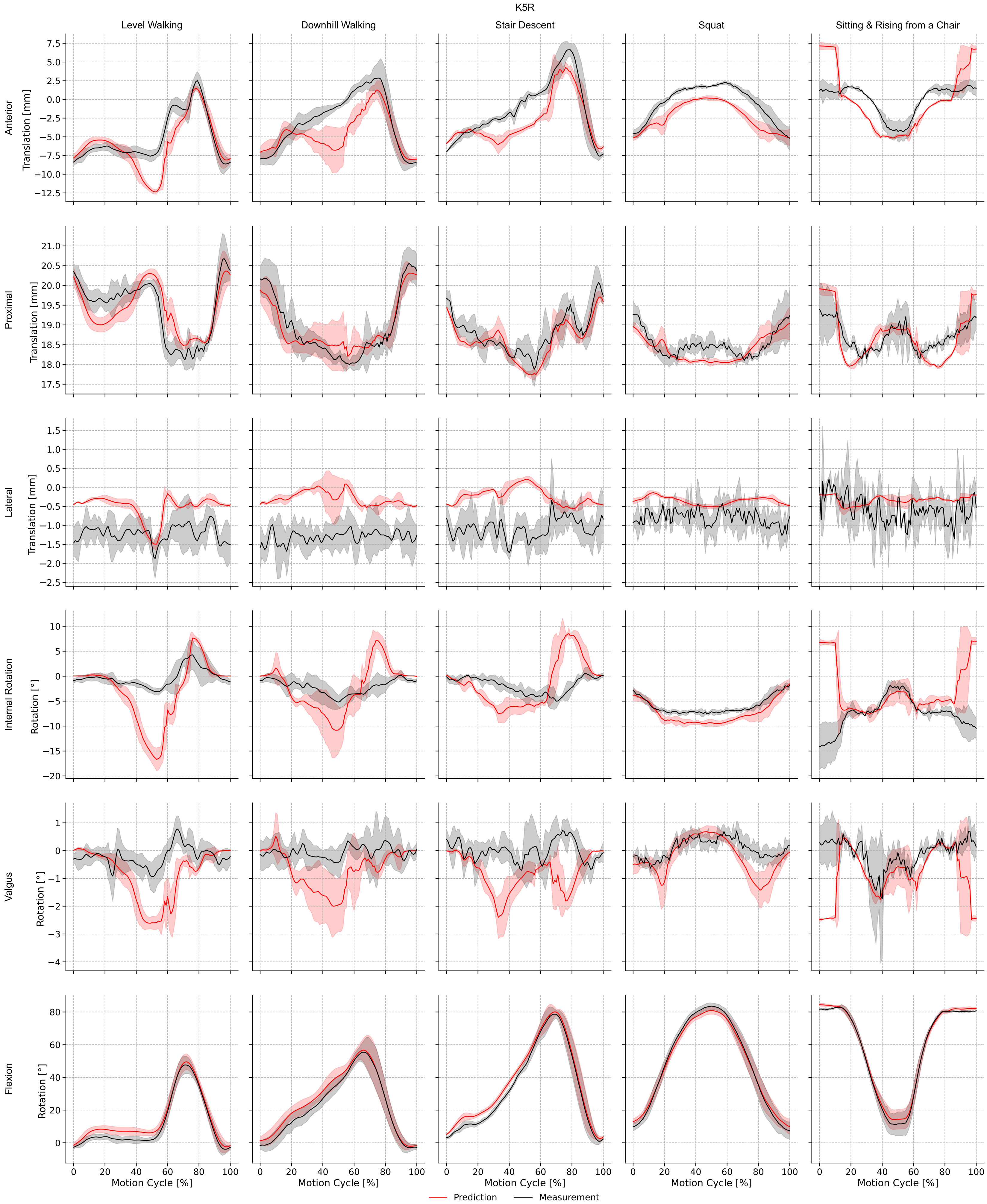

K5R
Level Walking
Downhill Walking
Stair Descent
Squat
Sitting & Rising from a Chair
Anterior
Translation [mm]
Proximal
Translation [mm]
Lateral
Translation [mm]
Internal Rotation
Rotation [°]
Valgus
Rotation [°]
Flexion
Rotation [°]
Motion Cycle [%]
Prediction
Measurement

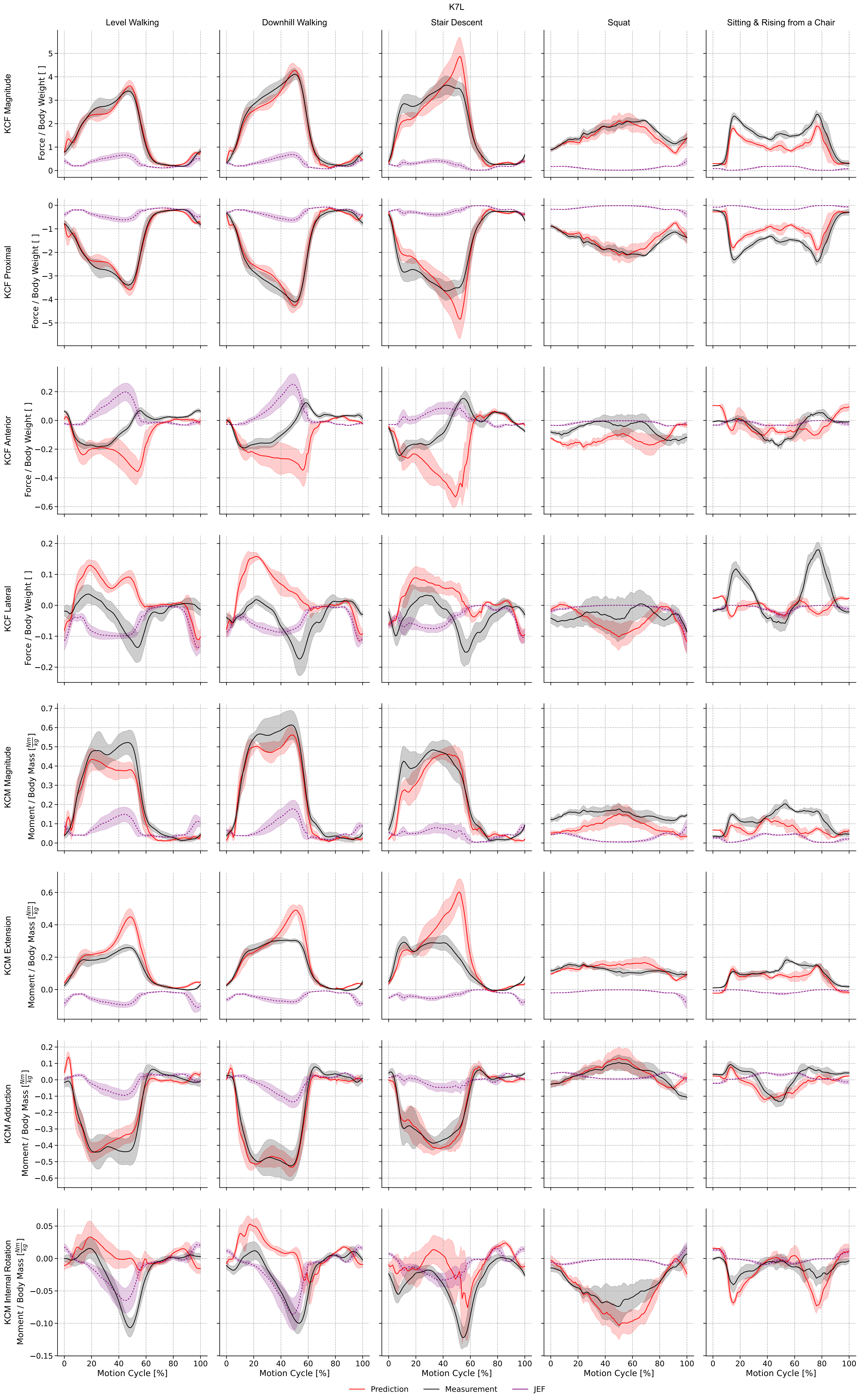
K7L
Level Walking
Downhill Walking
Stair Descent
Squat
Sitting & Rising from a Chair
KCF Magnitude
Force / Body Weight [ ]
KCF Proximal
Force / Body Weight [ ]
KCF Anterior
Force / Body Weight [ ]
KCF Lateral
Force / Body Weight [ ]
KCM Magnitude
Moment / Body Mass [Nm/kg]
KCM Extension
Moment / Body Mass [Nm/kg]
KCM Adduction
Moment / Body Mass [Nm/kg]
KCM Internal Rotation
Moment / Body Mass [Nm/kg]
Motion Cycle [%]
Prediction
Measurement
JEF

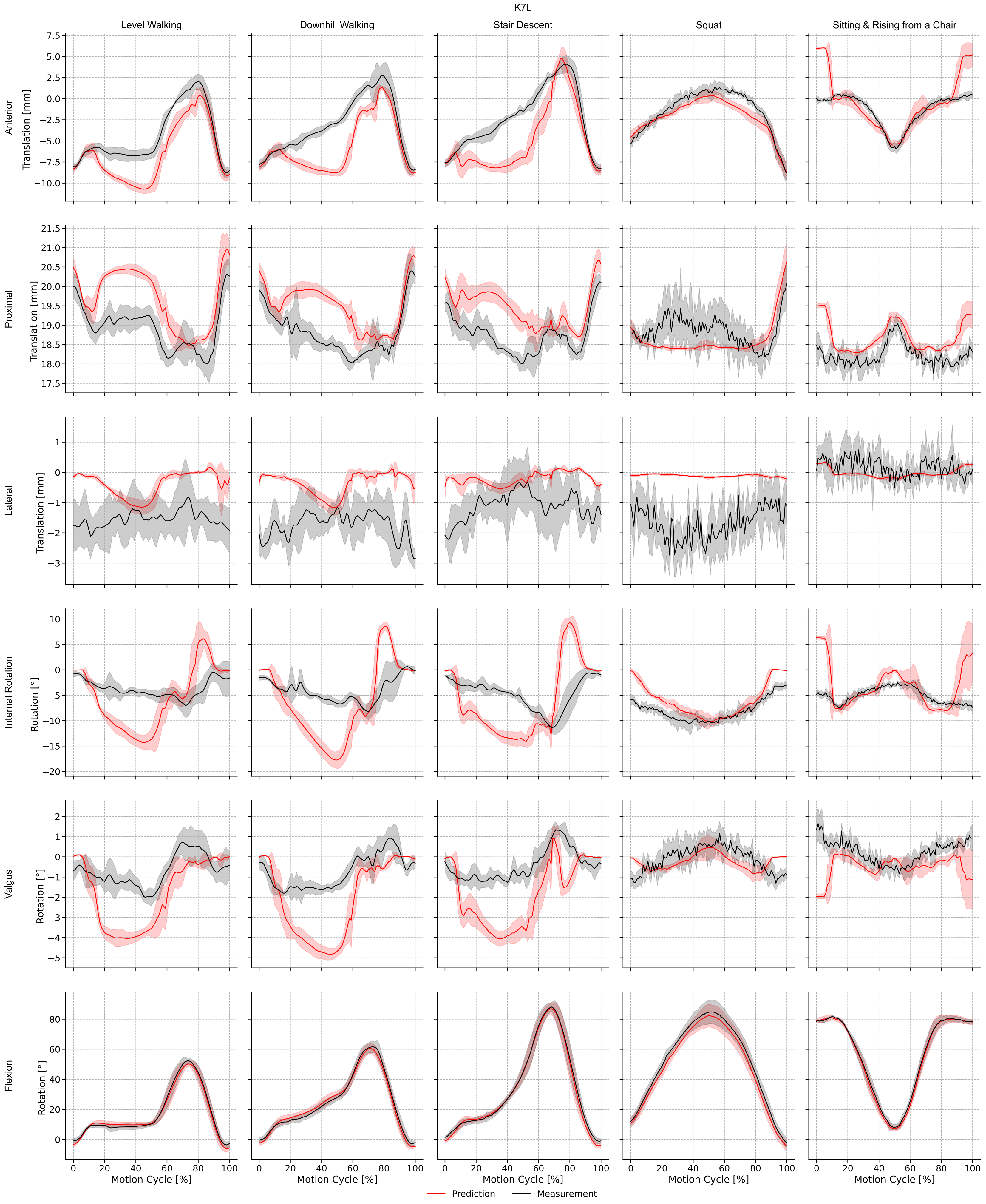

K7L
Level Walking
Downhill Walking
Stair Descent
Squat
Sitting & Rising from a Chair
Anterior
Translation [mm]
Proximal
Translation [mm]
Lateral
Translation [mm]
Internal Rotation
Rotation [°]
Valgus
Rotation [°]
Flexion
Rotation [°]
Motion Cycle [%]
Prediction
Measurement

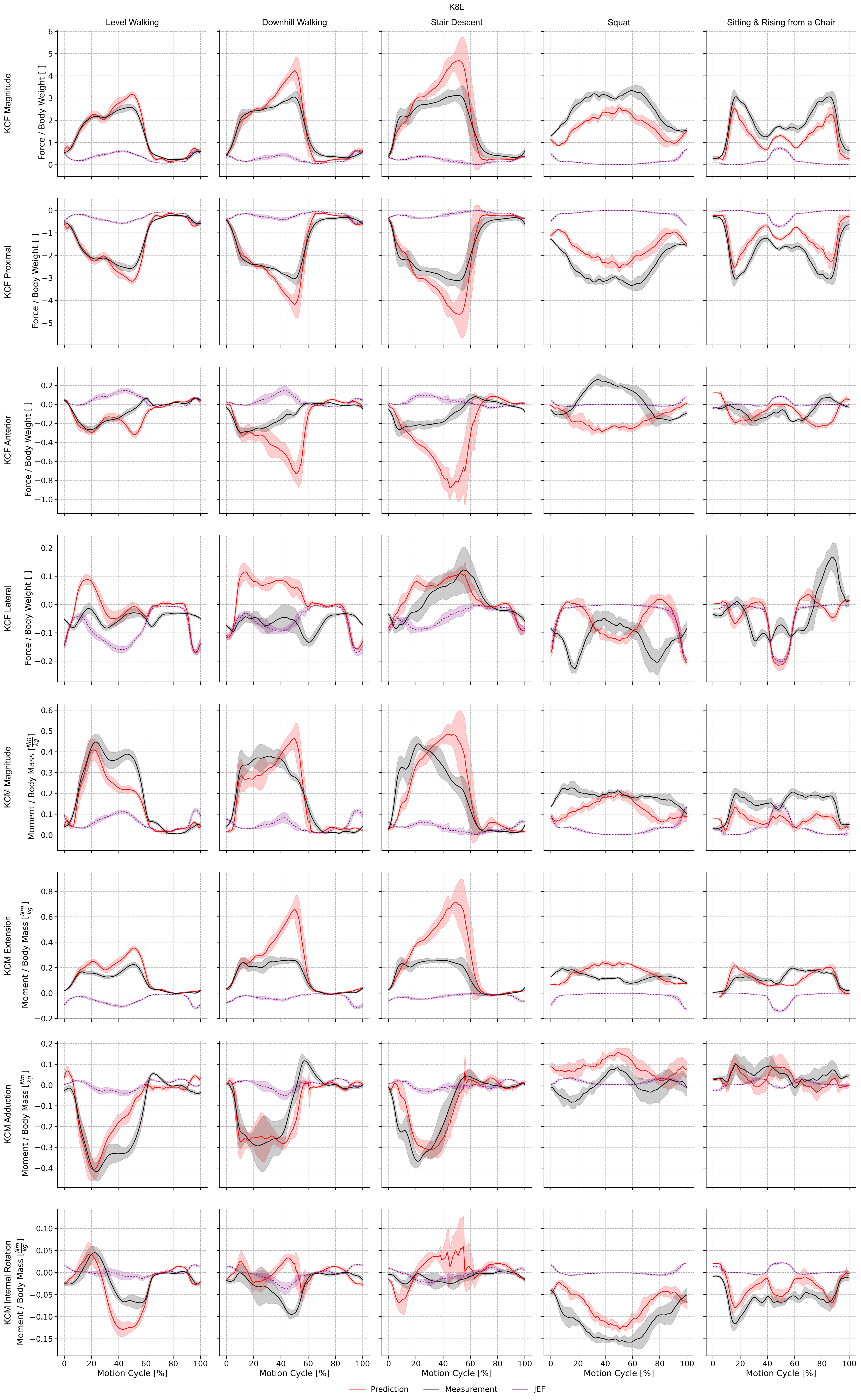

K8L
Level Walking
Downhill Walking
Stair Descent
Squat
Sitting & Rising from a Chair
KCF Magnitude
Force / Body Weight [ ]
KCF Proximal
KCF Anterior
KCF Lateral
KCM Magnitude
Moment / Body Mass [Nm/kg]
KCM Extension
KCM Adduction
KCM Internal Rotation
Motion Cycle [%]
Prediction
Measurement
JEF

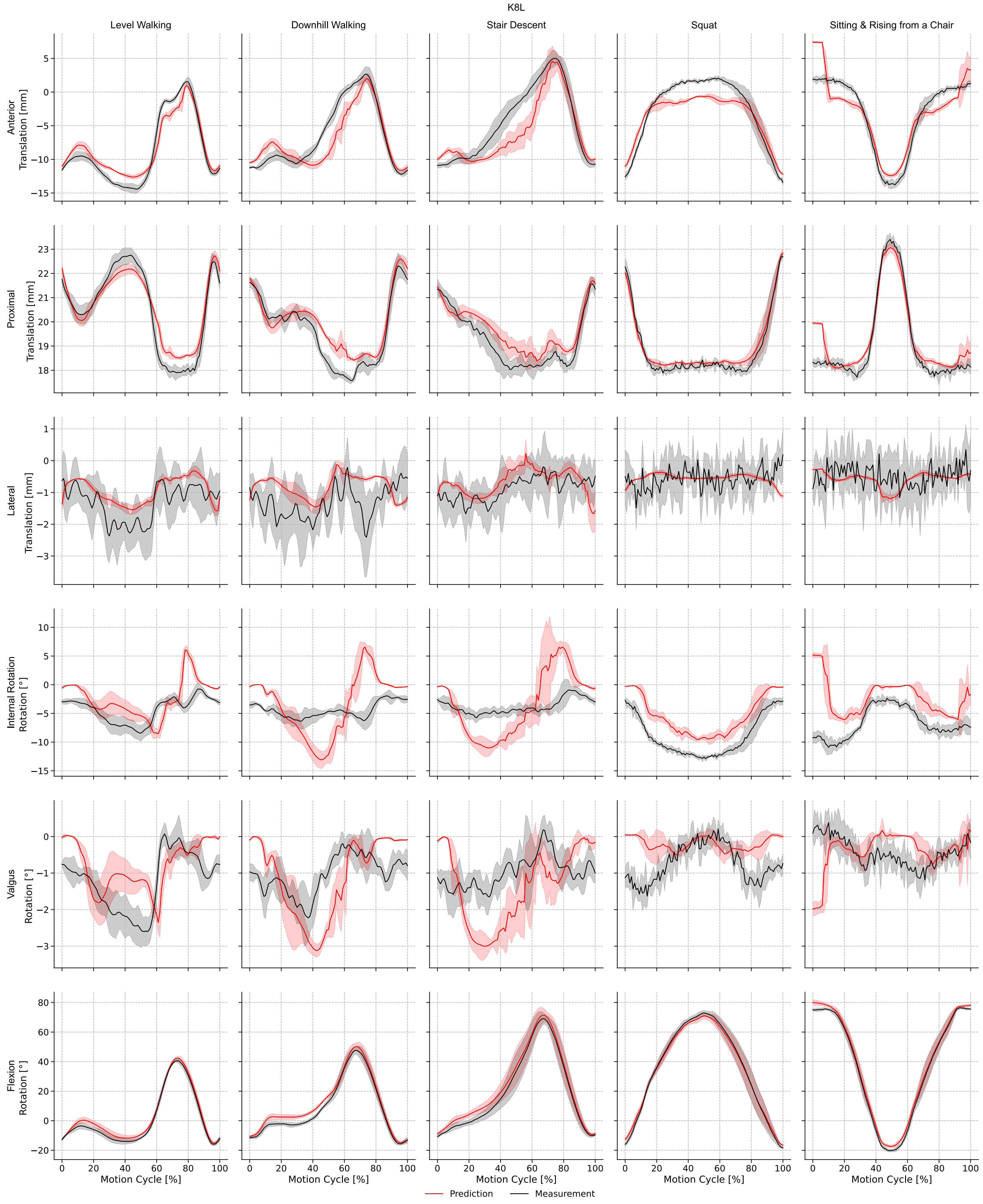
K8L
Level Walking
Downhill Walking
Stair Descent
Squat
Sitting & Rising from a Chair
Anterior Translation [mm]
Proximal Translation [mm]
Lateral Translation [mm]
Internal Rotation Rotation [°]
Valgus Rotation [°]
Flexion Rotation [°]
Motion Cycle [%]
Prediction
Measurement
